# High-accuracy hierarchical rRNA operon profiling resolves genomovar-level taxonomy and microdiversity using Nanopore sequencing

**DOI:** 10.64898/2026.06.01.729381

**Authors:** Kaiqin Bian, Ashwin S Sudarshan, Zijun Meng, Jinhao Yang, Megan Bachmann, Charles Bott, Katherine E. Graham, Ameet J. Pinto

## Abstract

Genomovar-level and intragenomic diversity cannot be resolved by conventional amplicon sequencing due to the limitation of fragment lengths and read accuracy, while the application of metagenomic profiling to a large number of samples can be resource intensive. Here, we report UltraRes-rrn, an integrated wet-lab and computational workflow for high-accuracy rrn (i.e., 16S– ITS–23S rRNA) operon profiling using Nanopore sequencing. By integrating ultra-long DNA recovery, long-read amplicon sequencing, and unique molecular identifiers (UMI)-based consensus correction, UltraRes-rrn obtains full-length 16S–ITS–23S rRNA operon consensus sequences with mean accuracies exceeding 99.98%. To achieve higher resolution, we propose a hierarchical rRNA operon profiling strategy in which concatenated 16S+23S rRNA genes (16S23S) serve as a primary and the internal transcribed spacer (ITS) provides a secondary marker. The 16S23S marker achieved discrimination at the genomovar level compared to either 16S or 23S rRNA genes alone, which mitigates the ITS-driven over-splitting observed with the full-length rrn operon and allows for larger proportion of data being classified at higher confidence thresholds. Further, ITS variation was strongly structured by tRNA occurrence patterns, suggesting that ITS can capture taxon microdiversity missed by either 16S or 23S rRNA gene sequences alone. The UltraRes-rrn workflow was applied to full-scale nitrogen removal reactors, revealing intraspecies diversity variation driven by different carbon regimes, which would not have been possible with a shorter gene sequence. In summary, UltraRes-rrn enables cost-effective community profiling at genomovar-level resolution in complex ecosystems.

## 1. Introduction

High resolution microbial community profiling (e.g., at intraspecies, intragenomic-level) can enhance our understanding of ecological functions and evolutionary dynamics of complex environmental communities (Karst et al., 2021; Leunda-Esnaola et al., 2024a; Woodruff et al., 2024). While metagenomic sequencing provides comprehensive genomic insights, it remains a labor-intensive, computationally demanding, and expensive method when the primary research objective is confined to taxonomic identification (Lema et al., 2023) and/or requires a large number of samples to be analyzed. Thus, amplicon sequencing remains the preferred high-throughput alternative for large-scale ecological surveys. Nanopore sequencing is a promising platform for cost-effective long-read sequencing, but its application for high-resolution taxonomy discrimination is hindered by its high sequencing error rate (∼2–5%) (Karst et al., 2021; Petrone et al., 2023). For instance, resolving intragenomic difference within the 4500 bp fragment of the full rRNA operon will require error rates substantially lower than 1% (Srinivas et al., 2025; Walsh et al., 2024). Unique Molecular Identifier (UMI)-based consensus sequencing has been proposed to address this limitation by assigning unique molecular identifiers to individual DNA template molecules before PCR amplification, allowing reads derived from the same original molecule to be clustered and collapsed into high-confidence consensus sequences (Deng et al., 2024; Karst et al., 2021; Lin et al., 2024). These UMI-based consensus sequencing strategies have been used to generate high-confidence consensus sequences (CCS) from reads with the same UMIs (Karst et al., 2021; Lin et al., 2024).

However, high-resolution community profiling cannot be achieved with improved accuracy alone. Conventional short-read amplicon sequencing targeting the V3-V4 region of the 16S rRNA gene resolves bacteria only to the genus or family level and cannot distinguish closely related species (Martínez-Porchas et al., 2016). Although the full-length 16S rRNA gene, supported by long-read sequencing platforms, improves taxonomic resolution, it still lacks sufficient discrimination capability between highly related species, such as *Escherichia coli* and *Shigella spp*., which share 99-99.8% 16S rRNA gene sequence identity (Bartoš et al., 2024; Devanga Ragupathi et al., 2017; Nygaard et al., 2020). The full-length rRNA operon (∼4500 bp), which captures additional sequence variation and phylogenetic signal spanning the 16S rRNA gene, internal transcribed spacer (ITS) region, and 23S rRNA gene, has emerged as a potential marker for higher resolution (Leunda-Esnaola et al., 2024b; Overgaard et al., 2024; Petrone et al., 2023). The ability to PCR amplify and sequence such a large amplicon also requires modification upstream of amplicon sequencing and data analyses, from DNA extraction to maximize unfragmented rrn operons available for PCR amplification to amplicon sequencing itself. For instance, conventional mechanical lysis methods, particularly high-energy bead-beating systems, over shear genomic DNA to fragment sizes below the length of the full rrn operon, thereby limiting recovery of intact templates for long-amplicon PCR (Bian et al., 2026; Knudsen et al., 2016). Moreover, successful amplification of these ultra-long targets requires high-fidelity polymerases, and super-high-accuracy basecalling and dedicated error-correction workflows to achieve accurate taxonomic profiling (Deng et al., 2024; Lin et al., 2024).

In this study, we report the development of UltraRes-rrn, an end-to-end framework that integrates ultra-long DNA extraction, UMI-based Nanopore amplicon sequencing, high-accuracy consensus sequence generation, and intragenomic diversity analysis of the markers in complex microbial samples. Instead of treating the full rRNA operon as a single marker, we propose a hierarchical utilization of the rrn operon regions, where species-level identification is performed via concatenated 16S+23S rRNA genes and ITS is utilized as a second marker to inform intraspecies and intragenomic diversity. By streamlining the process from wet-lab full rRNA operon amplicon library preparation to analysis pipeline, this study provides a robust workflow for high-accuracy microbial characterization beyond species-level resolution and improves amplicon sequence variant (ASV)-level profiling in complex ecosystems.

## 2. Materials and Methods

### 2.1. Evaluation of marker genes for prokaryotic taxonomy identification

A set of complete genome sequences of prokaryotes was retrieved from the *RefSeq* (release 212) and GTDB databases (release 207), as well as their taxonomic annotations (Chaumeil et al., 2022; O’Leary et al., 2016; Walsh et al., 2024). Sequences of 16S rRNA gene (16S), ITS region (ITS), 23 rRNA genes (23S), and 16S–ITS–23S rRNA operon (rrn) were extracted from complete genomes using ipcress from exonerate v2.4.0 with corresponding primer sets, length thresholds (**Table S1**), and parameters (-m 2 -M 64 -S 106 -p F -P) (Slater and Birney, 2005). Sequences of concatenated 16S+23S rRNA gene (16S23S) without ITS rRNA gene were generated by omitting ITS region from 16S–ITS–23S rRNA operon. In addition to regional comparisons, ITS sequences were further classified based on the tRNA gene occurrence patterns (i.e., presence and composition of tRNA genes), which were identified using tRNAscan-SE v2.0 under bacterial mode with default parameters (Chan et al., 2021). To minimize ambiguous calls, when a single region could be assigned to multiple tRNA occurrence patterns, we retained only the annotation with the highest tRNAscan-SE score. Average Nucleotide Identity (ANI) and pairwise identities were calculated for all complete prokaryotic genomes and different regions of the rrn operon, respectively. ANI values were determined between all pairs of genomes using skani (Shaw and Yu, 2023). To calculate pairwise identities for each region (16S, ITS, 23S, 16S23S, rrn, and ITS with different tRNA occurrence patterns), sequences of each region were split by genus and aligned independently using MAFFT v7 with the parameters --parttree --retree 2 --maxiterate 0 (Katoh et al., 2002). Pairwise identity was calculated from the aligned sequences using the EMBOSS tool distmat under the uncorrected p-distance model (-nucmethod 0) (Rice et al., 2000). Two taxonomy assignment matrices were used in this analysis. Phylogenetic genus, species, and genome assignments were determined using GTDB taxonomy (Conrad et al., 2024). Genomospecies (i.e., species-level group defined by whole-genome similarity) and genomovar (i.e., clusters of strains within a single species that are genomically distinct but cannot be differentiated by standard phenotypic tests) boundaries were defined using dual thresholds of ANI and alignment fraction (AF): >95% ANI with >60% AF for the same species, and >99.8% ANI with >85% AF for the same genomovar (Conrad et al., 2024; C. Jain et al., 2018; Rodriguez-R et al., 2023; Varghese et al., 2015). The correlation between genomic ANI and pairwise identity of each region was analyzed and visualized using R packages. Marker discrimination of closely related genomes or intragenomic sequences was further evaluated by receiver operating characteristic (ROC) analysis using pairwise identities of 16S, ITS, 23S, 16S23S, and rrn as predictors. Positive pairs were genomovar-level or same-genome and all others were negative. ROC curves were commutated using pROC in R (Robin et al., 2011).

### 2.2. *In silico* analysis of correlation between intact 16S–ITS–23S rRNA operon recovery and DNA fragment length

We performed *in silico* analysis using an ultra-long read dataset (SRR11027197) to quantify the relationship between DNA fragmentation and recovery of full-length rrn (M. Jain et al., 2018). The full-length rrn was extracted using ipcress from exonerate v2.4.0 with parameter -m 2 -M 64 -S 106 -p F -P (Slater and Birney, 2005). The full rrn recovery rates were then evaluated across increasing maximum read-length thresholds. At each threshold, recovery was calculated as the proportion of intact rrn-containing reads at or below that length relative to the total rrn copy number detected in the full dataset.

### 2.3. Evaluating DNA extraction protocols to maximize DNA yield and fragment length distribution

We tested range of protocols to optimize the recovery of high-molecular-weight (HMW) DNA while simultaneously ensuring high DNA yields so as to maximize the recovery of intact 16S– ITS–23S rRNA gene operon. To this end, we adapted two commercial extraction protocols, a mechanical lysis-based Qiagen DNeasy PowerWater Kit (PW, Qiagen, Germany) and an enzymatic-lysis based Wizard HMW DNA Extraction Kit (EN, Promega, USA), into two sequential extraction workflows, designated as DoubleBeads (DB) and EnzyBeads (EB), respectively (**Figure 1**). Specifically, the supernatant from the first lysis step from the standard protocol (PW or EN) was removed and the residual cell pellet was transferred to a PowerWater Bead Pro tube (Qiagen, Germany) and subjected to a second mechanical lysis step by vortexing for four minutes. The lysates from both rounds were merged prior to final purification. The detailed, step-by-step version of the protocols is available at protocols.io (https://dx.doi.org/10.17504/protocols.io.n2bvjkk8wgk5/v1). The four protocols were first tested on *Escherichia coli* (*E. coli*) cell cultures. *E. coli cell* cultures in mid-log phase were harvested by centrifugation (4,500 ×g, 2 min), washed in phosphate-buffered saline (PBS, VWR International, USA), and partitioned into 0.5 mL aliquots for the DNA extraction. Negative controls of UltraPure DNase/RNase-Free Distilled Water (Invitrogen, USA) were processed in parallel with the *E. coli cell* cultures. DNA yield for each protocol was quantified on the Qubit Flex fluorometer (Invitrogen, USA) using the 1× Qubit dsDNA High Sensitivity Assay Kit (Invitrogen, USA) following manufacturer instructions. The fragment size distribution was evaluated using a TapeStation 4200 system with the Genomic DNA ScreenTape Kit (Agilent Technologies, CA, USA). Data from the TapeStation were analyzed and visualized using the bioanalyzeR package. All DNA extracts were purified using ClenNGS (CleanNA, the Netherlands) at a 0.9× ratio following the manufacturer’s instructions.

**Figure 1.**
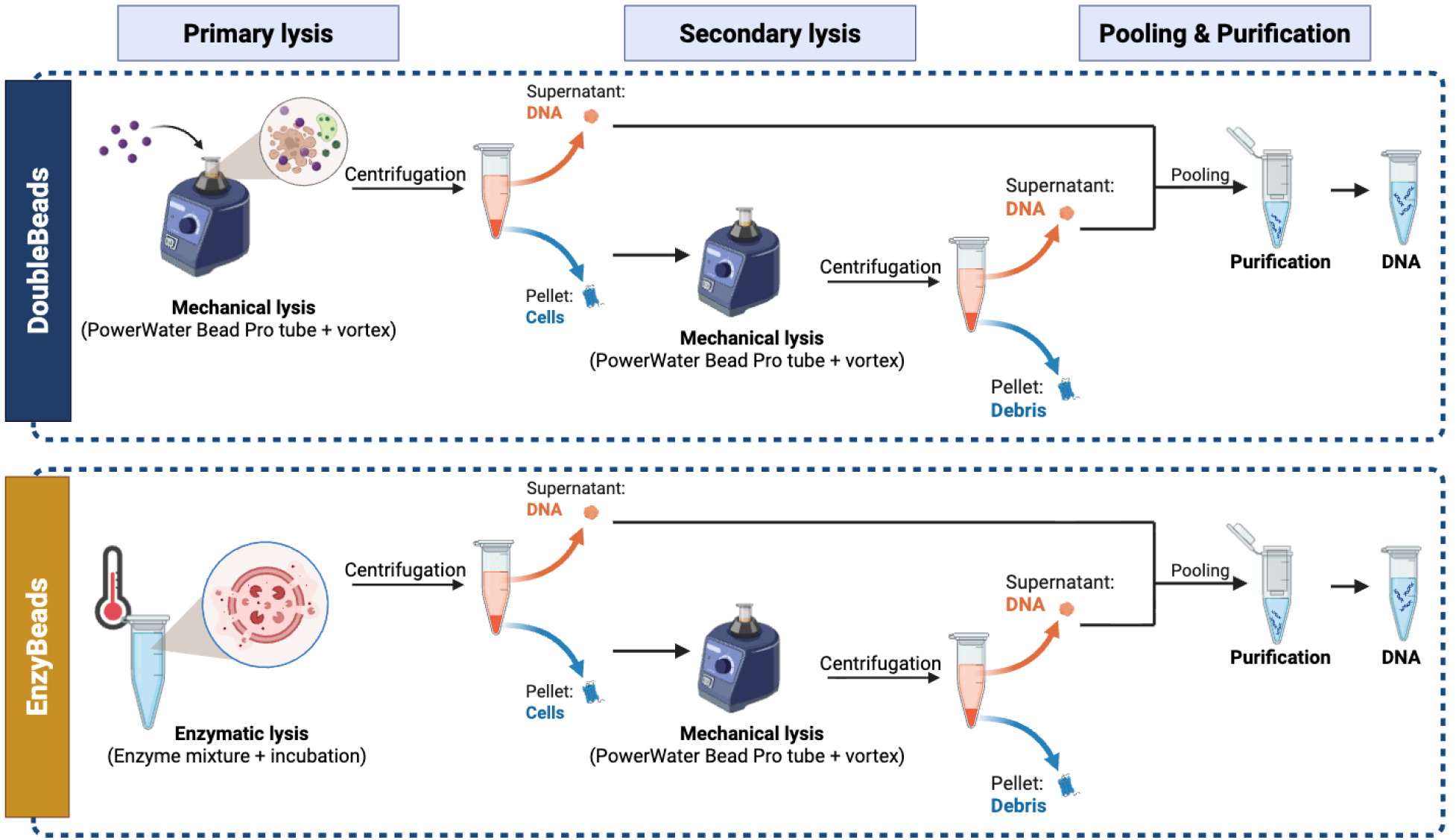
Schematic of the sequential DNA extraction procedures. The top panel illustrates the DoubleBeads (DB) method, which utilizes two mechanical lysis steps (PowerWater Bead Pro tube and vortex). The bottom panel shows the EnzyBeads (EB) method, which employs a primary enzymatic lysis (enzyme mixture with incubation) followed by a mechanical lysis. Both methods follow: primary lysis, centrifugation to separate free DNA (Supernatant) from intact cells (Pellet), secondary lysis of the cells from the pellet, another centrifugation to separate additional free DNA from debris, and pooling of both DNA-containing supernatants for DNA purification and elution.

### 2.4. Unique molecular identifier-based long-read Nanopore sequencing of full-length rRNA operon using a mock microbial community

DNA extracted from ZymoBIOMICS Gut Microbiome Standard (Zymo Research Corporation, USA) (Zymo Gut DNA) following the sequential ultra-long read recovery protocol optimized in section 3.2 was processed following the unique molecular identifier (UMI)-based long-read Nanopore sequencing workflow.

We first adapted a three-step strategy consisting of unique molecular identifier (UMI) tagging and two rounds of PCR amplification targeting 16S–ITS–23S rRNA operon (Karst et al., 2021; Lin et al., 2024). This approach was taken to facilitate read correction using a consensus-based approach. The primers and thermocycling conditions for tagging and amplification are listed in **Tables S2** and S3. The UMI tagging reaction was performed in a 20 μL volume containing 1x Platinum SuperFi II PCR Master Mix (Invitrogen, USA) and 200 nM each of the forward and reverse UMI tagging primers. The primer concentration was optimized to minimize non-specific amplification and primer-dimer formation for large amplicon targets. The thermocycling conditions for tagging included an initial denaturation at 94°C for 3 min, followed by 2 cycles of denaturation (94°C, 30 s), annealing (60°C, 15 s), and extension (68°C, 3 min) and concluding with a final extension at 68°C for 10 min. Following the tagging reaction, products were purified using CleanNGS magnetic beads at a 0.4× ratio. The beads were washed three times with 80% ethanol, air-dried for 5 min, and DNA was eluted in 20 μL of molecular biology-grade water. The purified UMI-tagged templates (8 μL) were subject to the first amplification round (PCR-1) in a 20 μL reaction. Each reaction contained 1x Platinum SuperFi II PCR Master Mix and 200 nM of each universal primer (UMI_16S_27F and UMI_23_U2428R, **Table S2**). The thermocycling program consisted of 25 cycles of denaturation (94°C, 15 s), annealing (60°C, 15 s), and extension (68°C, 3 min). PCR-1 products were purified using the same 0.4× CleanNGS bead protocol and eluted in 20 μL. To ensure sufficient DNA yield for Nanopore library preparation, a second amplification round (PCR-2) was conducted. A total of 5 μL or 15 μL of the purified PCR-1 product was used as the template for a 5-cycle PCR, utilizing the same thermocycling conditions and reagent concentrations as PCR1. The final amplicons were purified at a 0.4× bead ratio, eluted in 20 μL, and validated via gel electrophoresis before downstream sequencing.

Nanopore sequencing libraries of UMI-tagged 16S–ITS–23S rRNA amplicons were prepared using the Native Barcoding Kit (SQK-NBD114.24, Oxford Nanopore Technologies, UK) following the manufacturer’s optimized protocol for high-molecular-weight DNA. The final prepared libraries were quantified using a Qubit fluorometer to ensure optimal loading concentration. Sequencing was performed on MinION R10.4.1 flow cells for up to 72 hours to maximize data throughput. Raw sequencing signal data (POD5) were basecalled using Dorado v1.3.1 with the super-accuracy (SUP) model (dna_r10.4.1_e8.2_400bps_sup@v5.2.0) (nanoporetech/dorado). Low-quality (Q-score < 20) or out-of-range (length < 3500 bp or > 6000 bp) reads were removed. The remaining reads were demultiplexed and trimmed for barcodes and adapters for downstream processing.

### 2.5. High accuracy reads generation and error profiling

High-accuracy consensus sequences for the 16S–ITS–23S rRNA sequences were generated using a modified longread_umi framework (Karst et al., 2021; Lin et al., 2024). This pipeline was specifically optimized to handle the approximately 4.5 kb target PCR product length and the specific error profile of R10.4.1 chemistry. First, basecalled reads were quality filtered to remove reads with an expected error (EE) threshold exceeding 10% (Rognes et al., 2016) using VSEARCH (v2.15). To ensure the recovery of complete 16S–ITS–23S rRNA sequences and exclude fragmented templates or non-specific products, a strict length filter was implemented to retain only those sequences between 3,500 and 6,000 bp. Next, the 36-bp UMI pairs (18 bp at each terminus) were identified, and sequences were clustered into UMI bins based on shared UMI pairs, allowing for a maximum of 4 bp mismatches within the UMI sequences to account for potential stochastic sequencing errors. To provide sufficient statistical power for error correction and ensure the reliability of the resulting consensus, only UMI clusters containing a minimum of three independently sequenced reads (cluster size >= 3) were retained for the polishing stage. A multi-step iterative polishing workflow involving iterative polishing and taxonomic classification was executed to generate a high-fidelity consensus sequence for each UMI bin. An initial draft consensus was first produced and subsequently refined through three rounds of Racon (v1.4) polishing to correct structural and indel errors (Vaser et al., 2017). This was followed by final base-level polishing using one round of Medaka (v1.7), utilizing the specialized r1041_e82_sup model (https://github.com/nanoporetech/medaka). Minimap2 (v2.17) with the -ax map-ont flag was utilized to align raw reads to the consensus during each polishing iteration (Li, 2018). The final, high-accuracy UMI-consensus sequences (CCS) were then utilized as the basis for taxonomic classification and phylogenomic analysis. Sequences of 16S, ITS, 23S, 16S23S, rrn were extracted using ipcress with respective primers (**Table S1**), followed by primer trimming.

To enable error-rate quantification, two reference databases of amplicon sequence variants (ASVs) were constructed. rrn sequences were extracted with ipcress using the 16S–ITS–23S primer set (forward: 5′-AGRRTTYGATYHTDGYTYAG-3′; reverse: 5′-CCRAMCTGTCTCACGACG-3′) from (i) the ZymoBIOMICS Gut Microbiome Standard reference genomes (Zymo Research, USA) and (ii) a Q40-filtered PacBio HiFi metagenomic dataset (NCBI BioProject PRJNA680590). Extracted sequences were dereplicated with USEARCH v11 and denoised with UNOISE3 to generate the ASV references (Hunt et al., 2006; Klindworth et al., 2013). For each dataset, raw reads, consensus sequences, and ASVs were aligned to the corresponding reference database using minimap2 (-ax map-ont --cs). Secondary and supplementary alignments were then excluded with SAMtools view -F 2308. Error rates were calculated as the sum of mismatches, insertions, and deletions divided by alignment length. To construct a phylogenetic tree for each marker, pairwise alignments were first performed using a global alignment strategy with free end gaps. The phylogenetic tree was subsequently built using the Neighbor-Joining (NJ) method. Evolutionary distances were computed using the Tamura-Nei genetic distance model, and the final tree was generated without an outgroup (Janssen et al., 2018). Silhouette analysis was conducted to evaluate phylogenetic clustering performance (Rousseeuw, 1987). For the phylogenetic tree of each region, pairwise patristic distances were computed using ape R package and clustered by average linkage hierarchical clustering (UPGMA). Cluster solutions were further evaluated across k = 2–10, and the optimal solution for each region was selected based on the highest mean silhouette width. For each region, tip-level silhouette widths were summarized by the mean, median, standard deviation, minimum, and maximum values to quantify the distribution of clustering quality in terms of within-cluster cohesion and separation from the nearest alternative cluster.

### 2.6. Environmental sample collection, UMI-based sequencing, and data processing

Plastic biofilm carriers were collected from two full-scale Integrated Fixed-film Activated Sludge (IFAS) partial denitrification-anammox (PdNA) reactors (IFAS 1 and IFAS 2) at the Hampton Roads Sanitation District James River Treatment Plant (JRTP) from February to December 2023. A detailed overview of JRTP facility has been described in previous research (Bachmann et al., 2025). In brief, IFAS1 and IFAS2 are integrated PdNA reactors in the second anoxic zone of a five-stage Bardenpho process, enhancing JRTP’s biological nitrogen removal. The reactors were operated in parallel under different carbon source regimes to evaluate the process stability, nitrogen removal efficiency, and potential for cost optimization under full-scale conditions. IFAS 1 received external carbon (glycerol) in addition to step-fed primary clarifier effluent (PCE), whereas IFAS 2 relied solely on step-fed PCE as the carbon source for partial denitrification, representing an economic but naturally carbon-limited regime. The information on the sampling date and corresponding operational phases is provided in **Table S4**.

To ensure the samples represented a biological steady state, collection was only performed when the plant’s Distributed Control System trends indicated stable influent flow and consistent NH□□ and NOx concentrations in the upstream aerobic effluent. Upon each collection, the plastic media were immediately submerged in RNAlater solution (Invitrogen, Waltham, MA) to stabilize the nucleic acids and shipped on ice to Georgia Institute of Technology. Prior to DNA extraction, plastic biofilm carriers of each sample were immersed in a 50 mL sterile centrifuge tube with 40 mL 1X phosphate-buffered saline (PBS), followed by vortexing for 5 minutes in order to dislodge attached biofilms. The agitated IFAS media solution was then concentrated by centrifugation at 4000 rpm for 10 minutes. After the media and supernatant were removed, the pellet was washed twice by resuspending in 40 mL 1X PBS and centrifuging at 4000 rpm for 5 min. The supernatant was discarded after each wash to remove residual enzyme inhibitors. The final sediments were subjected to the sequential ultra-long DNA extraction and purification protocol optimized in section 3.2. Finally, the DNA extracts were diluted 10-fold to mitigate the effects of any inhibitors for downstream UMI-based Nanopore sequencing described in section 2.4. High-accuracy consensus sequences were generated following the pipeline described above (section 2.5) for downstream taxonomic, ecological, and statistical analyses.

### 2.7. Bioinformatics and statistical analysis for mock community and environmental samples

As the full-length amplicons spanned the 16S–ITS–23S rRNA operon, region-specific datasets were generated from each consensus sequence for comparative analysis across marker regions using ipcress with primers for each region (**Table S1**). The analyzed regions included the 16S, ITS, 23S, 16S23S, rrn. Sequences of each region were classified independently using custom RDP classifiers, and taxonomic assignments were obtained from domain to species level with a confidence score threshold of > 0.8 (Wang et al., 2007). The RDP classifiers were trained on region-specific sequences and taxonomy annotation derived from *RefSeq* and GTDB datasets constructed in section 2.1. To evaluate classification consistency across marker regions, read-level taxonomic concordance and discordance were assessed by comparing taxonomic assignments from each region to those obtained from the 16S23S region. ANCOM-BC2 was implemented using region-specific count tables for differential abundance analysis of microbial taxa identified via different marker regions (Lin and Peddada, 2024). Prior to applying the confidence score cutoff, classifier confidence score distributions were also compared across regions to assess differences in assignment certainty. Alpha diversity was quantified using the Shannon index, while beta diversity was evaluated via Bray–Curtis dissimilarity and visualized using Principal Coordinates Analysis (PCoA). Core taxa were defined separately for each treatment group as taxa detected at a relative abundance of at least 0.1% in at least 75% of samples within that group. Core taxa were further classified as shared core, Glycerol+PCE-specific core, or PCE-only-specific core according to their group-level prevalence patterns.

## 3. Results and discussion

### 3.1. Hierarchical utilization of the 16S–ITS–23S rRNA operon captures genomovar-level resolution and intragenomic heterogeneity

We compared pairwise identity of sequences of each marker (i.e., 16S, ITS, 23S, 16S23S, and rrn) of all genomes collected from *RefSeq* and GTDB against their whole-genome pairwise ANI to assess the concordance between marker-level and genome-wide divergence. Analysis encompassed 47,998 complete genomes from which we extracted 252,765 16S rRNA genes, 244,588 ITS regions, 244,665 23S rRNA genes, 244,432 concatenated 16S+23S rRNA gene sequences, and 252,227 full-length 16S–ITS–23S rRNA operon. Length distribution and pairwise identity of each region are shown in **Figures S1** and **S2**. While GTDB provides baseline annotations of each genome (e.g., genus, species, assembly accession), it lacks the resolution to differentiate intra-species diversity (Walsh et al., 2024). Given the high concordance between GTDB- and ANI-based taxonomic assignment at the species level (**Figure S3**), we further applied whole-genome ANI and AF thresholds to resolve GTDB-assigned species into distinct genomovars (C. Jain et al., 2018).

As shown in **Figures 2A** and **2C**, both 16S and 23S rRNA genes, which are widely used as phylogenetic markers for genus- and species-level taxonomic identification, remain identical at the same GTDB-species level, with a high-density hotspot above 99.9% identity even among genome pairs with ANI below 99.8%, indicating a decoupling between rRNA gene divergence and genome-wide divergence (Rodriguez-R et al., 2023; Woodruff et al., 2024). The conservation of 16S and 23S rRNA genes results in it being unlikely to distinguish genetically divergent strains within a species. Conversely, ITS pairwise identity spans a broad range (∼80%–100%) among genome pairs exceeding 95% ANI (**Figures 2B and S4B**). In addition, ITS region exhibited the greatest length variability across the database compared to the highly constrained regions (i.e., 16S and 23S rRNA genes), resulting in the wider length distribution of 16S–ITS– 23S rRNA operon (**Figure S1**). This observation is consistent with the lower functional constraint and faster evolutionary rate of ITS relative to 16S and 23S rRNA genes (Stewart and Cavanaugh, 2007a). Although ITS has been utilized as a genetic marker for taxonomic identification and genotyping of specific bacterial isolates, such as *E. coli* and *Bacillus cereus*, identical ITS from different subspecies or divergent ITS from the same genome were both observed in the comprehensive database (**Figure 2B**), regardless of whether classifications were defined based on whole-genome ANI thresholds or GTDB taxonomy (Asthana et al., 2025; Badi et al., 2021; Hu et al., 2022; Ruegger et al., 2014). Thus, its universal application in taxonomic identification remains limited in culture-independent environmental samples with complex microbial composition.

**Figure 2.**
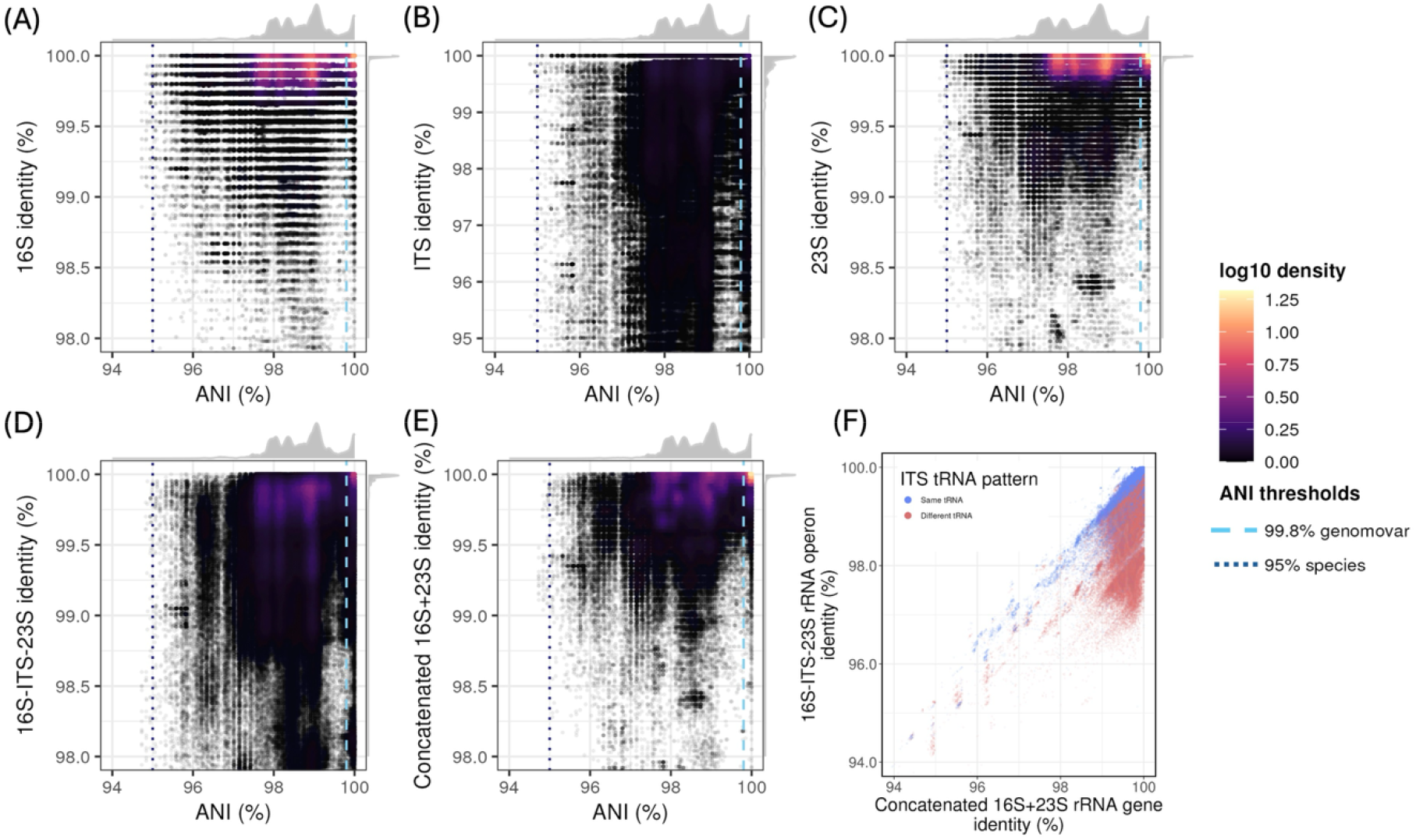
Correlation analysis between whole-genome Average Nucleotide Identity (ANI) and intraspecific pairwise sequence identity across different rRNA regions for genomes from the same species. Each panel displays a density plot comparing ANI with pairwise identity of: (A) 16S, (B) ITS, (C) 23S, (D) 16S–ITS–23S rRNA operon, and (E) concatenated 16S+23S rRNA genes. (F) Comparison of the pairwise identity between the 16S-ITS-23S rRNA operon and concatenated 16S+23S rRNA genes from same species. In panels (A-E), the color gradient represents the log10-transformed density of data points. Top and right marginal density curves (solid grey) represent the univariate distributions of ANI and region-specific identity, respectively. The color gradient represents the density of data points at each position. Vertical dashed lines indicate whole-genome ANI thresholds of 95.0% and 99.8%, corresponding to commonly used species- and genomovar-level genomic boundaries, respectively. In panel F, colors indicate whether each sequence pair shares the same or different tRNA occurrence patterns.

To overcome the limitations of individual marker, we evaluated the performance of the full 16S– ITS–23S rRNA operon and concatenated 16S+23S rRNA genes. By combining slow (16S and 23S rRNA genes) and rapidly (ITS) evolving regions, the full 16S–ITS–23S rRNA operon maximizes information content and improves taxonomic resolution over individual regions (Won et al., 2024). However, the full 16S–ITS–23S rRNA operon exhibited a greater pairwise identity variance than the 16S or 23S rRNA genes alone (**Figures 2D and S4**) (Stewart and Cavanaugh, 2007a). A substantial fraction of intraspecies sequence pairs exhibited below 98% 16S–ITS–23S rRNA operon identity (**Figures 2D and S4D**). Notably, concatenated 16S+23S rRNA genes (**Figure 2E**) yields more concentrated identity distributions at the genomovar level than either region alone or full 16S–ITS–23S rRNA operon (**Figures 2A-D**). The full 16S–ITS–23S rRNA operon shows a long tail extending to lower identity compared to concatenated 16S+23S rRNA genes, indicating its greater sequence variability (**Figure 2F**). This observation confirmed that the greater variance of the full 16S–ITS–23S rRNA operon is primarily driven by the heterogeneous evolutionary dynamics of the ITS region (Stewart and Cavanaugh, 2007b; Sun et al., 2013a; Won et al., 2024). We further evaluated the resolution of different markers at the genomovar level and intragenomic sequences using receiver operating characteristic (ROC) analysis (**Figure 3**). The area under the ROC curve (AUC) of the concatenated 16S+23S rRNA is 0.845 and 0.912 for genomovar and same-genome level, respectively, exhibiting the highest discriminatory ability among all tested markers (**Figure 3**). The concatenated 16S+23S rRNA gene leverages phylogenetically informative sites of both conserved subunits while excluding the hypervariable ITS region, which lacks correlation with taxonomic relatedness. Therefore, the 16S23S marker enhances taxonomic identification sensitivity to the genomovar level and avoids overestimation of microbial diversity (Pan et al., 2023; Sun et al., 2013b).

**Figure 3.**
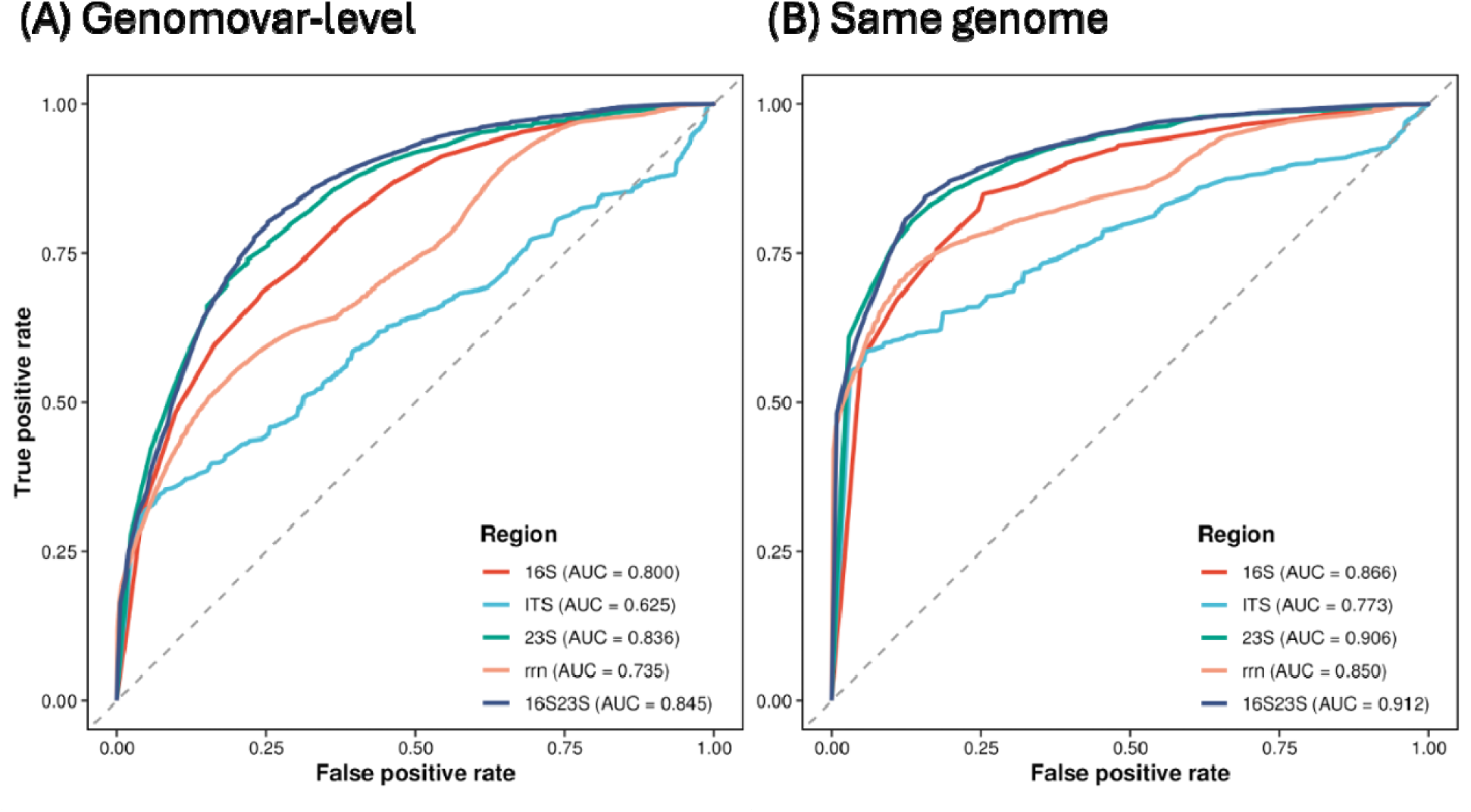
Receiver operating characteristic (ROC) curves comparing the ability of identities from the 16S rRNA gene (16S), internal transcribed spacer (ITS), 23S rRNA gene (23S), concatenated 16S+23S rRNA genes (16S23S), and full 16S–ITS–23S rRNA operon (rrn) to distinguish (A) genomovar-level pairs and (B) same-genome pairs. The area under the ROC curve (AUC) represents the overall discriminatory ability of each marker across all identity thresholds.

Despite its heterogeneity, the ITS region is structured by highly conserved transfer RNA (tRNA) genes (Leunda-Esnaola et al., 2024b; Stewart and Cavanaugh, 2007b). Among all extracted ITS sequences, 71.6% (175,152 out of 244,588) carried at least one tRNA gene (**Text S1**), indicating that tRNA gene insertion is widespread. We observed 104 distinct patterns, accounting for the combination of tRNA type and copy number of each type. The combination of Ala+Ile was the most dominant (103,130), followed by Glu-only (52,450), Ala-only (12,985), and Ile-only (3,201). To further assess the impact of tRNA gene occurrence patterns on the sequence divergence, we performed pairwise comparisons of all sequences within the same genus for each marker, linked the pairwise identity to their ANI, and stratified them by the tRNA gene occurrence pattern concordance (**Figure 4**). For each of the 16S, 23S, and 16S23S markers, identity distributions were nearly identical regardless of tRNA pattern concordance (**Figure S5A, S5B, and 4B**), indicating that these conserved regions are decoupled from ITS structural variation. In contrast, ITS sequences sharing the same tRNA gene occurrence patterns exhibited higher identity than those with different patterns at both interspecies (same genus but not same species) and intraspecies (same species) levels (**Figures 4A and S6**), with two distinct peaks emerging within same-genus/different-species and same-species/different-genome comparisons. Similarly, full 16S–ITS–23S rRNA operon identities differed between sequence pairs with the same and different tRNA occurrence patterns (**Figure 4C**). Direct comparison with 16S23S identities showed that the tRNA gene occurrence pattern was a strong indicator of ITS-driven divergence. Pairs sharing the same tRNA pattern showed agreement between rrn and 16S23S identities, whereas pairs with different tRNA patterns exhibited lower and more dispersed full-operon identities despite high 16S23S similarity (**Figure S7**). This decoupling persisted even among rrn from the same genome, indicating that ITS structural heterogeneity can dissociate rrn identity from genome-level relatedness. Thus, incorporating the ITS region increases the risk of misclassifying rRNA operons originating from a single species into multiple distinct species, which is compounded by the limitations of current reference databases. The incomplete Metagenome-Assembled Genomes (MAGs) frequently fail to capture the full spectrum of unique ITS sequences between the highly conserved 16S and 23S regions. As the phylogenetically informative signal is retained in the conserved 16S and 23S and the additional divergence arises from ITS-associated heterogeneity, the 16S23S marker is more stable for taxonomic identification, whereas the ITS tRNA-pattern variation captures a secondary layer of intragenomic and intraspecific operon heterogeneity.

**Figure 4.**
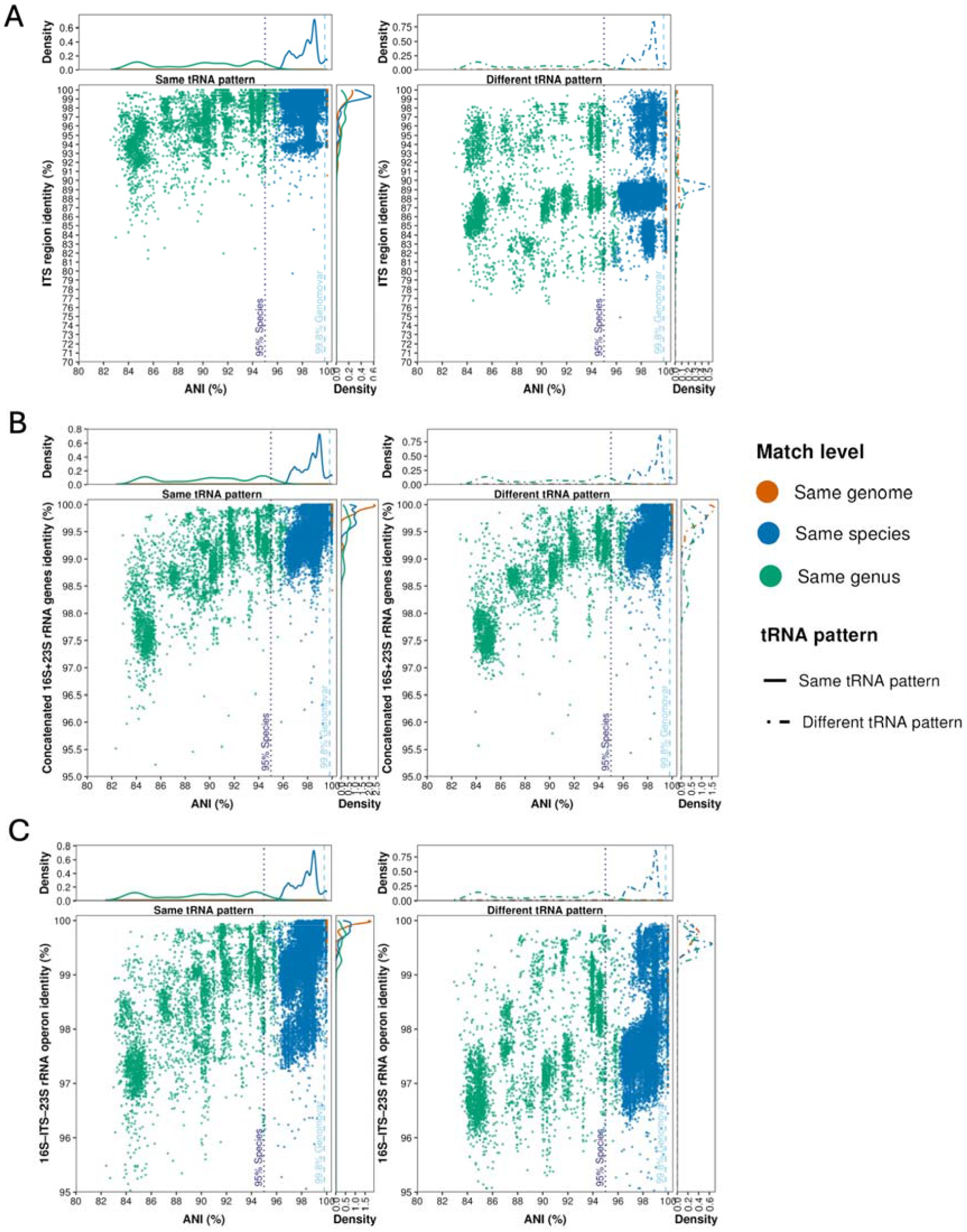
Density plots of whole-genome ANI versus pairwise sequence identity across rRNA regions, stratified by tRNA gene occurrence pattern concordance. (A) ITS, (B) concatenated 16S+23S rRNA gene, and (C) 16S–ITS–23S rRNA operon. In each panel, the left and right plots represent the sequence comparisons with the same or different tRNA occurrence patterns, respectively. The top and right marginal density curves show the univariate distribution of ANI and region-specific identity, respectively. Point colors in the main scatterplots and line colors in the marginal density plots indicate the taxonomic relationship of each sequence comparison, as defined by GTDB taxonomy. Vertical dashed lines indicate whole-genome ANI thresholds of 95.0% and 99.8%, corresponding to commonly used species- and genomovar-level genomic boundaries, respectively.

While divergence of ITS region weakens the concordance between full 16S–ITS–23S rRNA operon identity and taxonomic relatedness, the ITS region provides insights into intragenomic diversity, which underpins the higher ecological adaptation potential of prokaryotes (Saraf et al., 2024; Stadthagen-Gomez et al., 2008). The heterogeneity within the ITS is thus driven by functional requirements rather than taxonomic speciation. It was also reported that the rRNA operon diversity is primarily generated by homologous recombination and replication slippage, which induce significant cross-operon nucleotide exchanges, rather than simple point mutations (Gürtler, 1999; Sultanov and Hochwagen, 2022). This interplay of recombination and mutation explains how bacteria maintain the concerted evolution of ribosomal genes while actively generating structural divergence. Previous studies have illustrated that intracellular tRNA gene pools are closely linked to environmental conditions (e.g., temperature, nutrition levels) and ITS with different tRNA genes enables prokaryotes to differentially regulate the expression of specific rRNA operons in response to fluctuating environmental pressures or across distinct growth stages (Fruchard et al., 2025; Jain and Cope, 2024; Tiefenbacher et al., 2024). Based on these observations, we propose a hierarchical framework for multi-resolution rRNA operon profiling. First, the concatenated 16S+23S sequence serves as the primary marker for genomovar taxonomic identification. Second, the ITS region with its tRNA gene occurrence patterns is leveraged to infer intragenomic diversity and potential ecological variation. In this secondary analysis, ITS sequences should first be grouped according to the tRNA gene occurrence pattern, and the diversity can be assessed both within the same pattern ITS sequences and across the rRNA operons within the genomovar.

### 3.2 Sequential DNA extraction strategy of ultra-long DNA recovery for rRNA operon

Maximizing DNA recovery is necessary to ensure sequencing data is representative of the microbial community, while ensuring DNA integrity is required to prevent target fragmentation of the rrn operon and enable efficient capture of the near-complete rrn operon during UMI tagging and subsequent PCR amplification. Unlike short-read amplicons (e.g., 16S rRNA gene V3-V4 region, ∼450 bp) or full-length 16S (∼1.5 kb), the longer length (∼4.5 kb) of 16S–ITS– 23S rRNA amplicon product necessitated higher requirements for the integrity of DNA fragments. Although previous research demonstrated that longer templates improve the capture of targets, the critical fragment length required for rRNA operon recovery has not been systematically quantified (Manzari et al., 2020). To address this knowledge gap, we performed in silico analysis to evaluate the recovery rate of 16S–ITS–23S rRNA operons as a function of DNA fragment sizes. As shown in **Figure 5A**, the likelihood of fragmentation within the 16S– ITS–23S rRNA operon is inversely proportional to the DNA fragment length. Specifically, fragments shorter than 18 kb resulted in the fragmentation of 75% of the 16S–ITS–23S rRNA operon. The recovery curve displays a sharp inflection point around ∼52 kbp, below which the recovery rate declines precipitously. Thus, a fragment length >52 kb is essential to achieve a template recovery rate exceeding 75% and a fragment length >118 kb is required to recover 95% 16S–ITS–23S rRNA operon targets. These results underscore that preserving ultra-long DNA is not merely an ideal for sequencing quality, but also a critical requirement to maximize capture and subsequent sequencing of the rRNA operon spanning 16S–ITS–23S rRNA regions.

**Figure 5.**
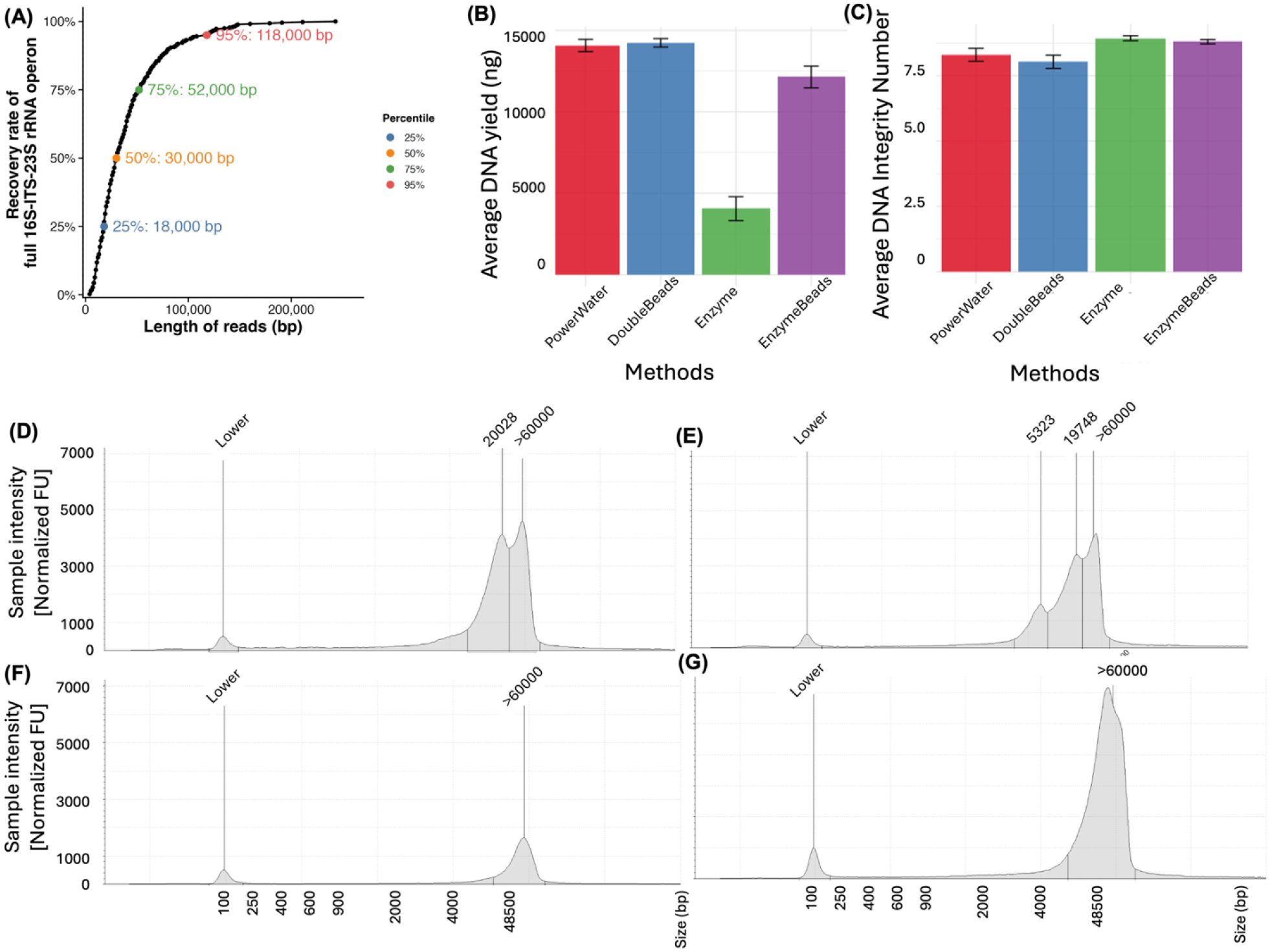
(A) Correlation between fragment size and recovery rate of intact 16S–ITS–23S rRNA operon. (B) DNA yield and (C) DNA integrity of different DNA extraction methods. Fragment distribution of different extraction methods. (D) PowerWater (PW), (E) DoubleBeads (DB), (F) Wizard HMW DNA Extraction (EN), and (G) EnzyBeads (EB). Peaks annotated with Lower indicate the Lower Marker in the TapeStation analysis.

However, conventional extraction protocols based on standard mechanical and enzymatic lysis failed to meet these requirements. While mechanical lysis methods (e.g., bead beating) ensures efficient cell disruption, they frequently induce DNA over shearing, leading to fragmentation of 16S–ITS–23S rRNA operon. In contrast, gentler enzymatic lysis approaches reduce DNA fragmentation but may result in incomplete lysis of some taxa (e.g., gram-negative bacteria), resulting in systematic underrepresentation and biased taxonomic profiles (Albertsen et al., 2015; Pinzauti et al., 2022). To balance the cellular lysis with DNA integrity, we tested two successive extraction strategies (DB and EB) that utilized a secondary mechanical lysis step to lyse any remaining intact cells for two representative mechanical (PW) and enzymatic (EN) extraction protocols, respectively. We compared the DNA yield and integrity across the four methods. The DNA integrity number (DIN), a key metrics for DNA degradation, was selected as the primary indicator for evaluating the structural integrity of the extracted DNA (Hiramatsu et al., 2023). As shown in **Figures 5B and 5C**, the successive strategies (DB and EB) yielded higher amounts of DNA as compared to the original single-step PW and EN methods, respectively. Thus, the secondary lysis recovered additional DNA from the cell pellets collected post primary lysis. However, in addition to the two shared fragment peaks at approximately 20 kb and 60 kb observed in both PW and DB, the secondary mechanical lysis in the DB protocol generated an additional peak at approximately 5323 bp (**Figures 5D and 5E**), suggesting that twice mechanical lysis resulted in DNA over-shearing. In contrast to the PW and DB methods, both EN and EB yielded DNA with higher DIN and fragment size exceeding 60 kb (**Figures 5F and 5G**). This peak size aligns with the requirement identified in silico for recovery of a representative template pool for the 16S–ITS–23S rRNA operon. In addition, the EB strategy achieved higher DNA yield (12135 ± 666.78 ng) compared to EN (4050 ± 733.72 ng), which are also comparable to DNA yield of DB (14230 ± 259.90 ng), indicating that the secondary lysis following the enzymatic lysis successfully recovered DNA from lysis-resistant cells without compromising the structural integrity. Therefore, the enzymatic-based successive DNA extraction method, EB, is identified as the most optimal method for the long-amplicon sequencing.

### 3.3. Coupling concatenated 16S+23S rRNA genes with UMI-based error correction enables genomovar-level microbial profiling in a mock microbial community

After establishing an optimized DNA extraction workflow, we further combined ultra-long read recovery and multi-resolution rRNA operon profiling with UMI-based consensus sequencing to reduce the intrinsic error rate of Nanopore sequencing. This integrated workflow, termed ULTRA-MultiRes-RRN, was then applied to the ZymoBIOMICS Gut Microbiome Standard. Before UMI-based polishing, the raw Nanopore reads had a mean accuracy of 98.6%.

We generated a total of 15288 consensus sequences (CCS) of full 16S–ITS–23S rRNA operon with a mean accuracy exceeding 99.8% and 99.98% against the ZYMO- and PacBio-derived references, respectively (**Figure 6A**). The four theoretically lowest-abundance species were *Clostridium perfringens, Salmonella enterica, Enterococcus faecalis*, and *Akkermansia muciniphila. C. perfringens* and *S. enterica* were undetected in both our Nanopore and PacBio datasets, whereas *E. faecalis* and *A. muciniphila* were absent only from the PacBio dataset. The absence of *A. muciniphila*, despite a relative abundance of 0.87%, suggests that PacBio detection was further limited by fragmented input reads, with Q40-filtered metagenomic reads showing a mean length of 7.3 kb, and a peak size of 4.7 kb. Residual errors were enriched in homopolymer regions, consistent with limitations of Nanopore sequencing technologies (**Figure S8**) (Overgaard et al., 2024). Error profiles of CCS for the other regions are shown in **Figure S9**. As the PacBio reference was constructed directly from Q40-filtered HiFi reads, it avoids the assembly and potentially PCR-associated errors. In contrast, the Zymo reference was derived from hybrid genome assemblies, which may retain residual assembly errors as previously reported (Karst et al., 2021; Lin et al., 2024). Thus, the PacBio reference provides a more robust benchmark for CCS accuracy. The higher identity to this PacBio-derived reference indicates the high accuracy of our CCS dataset. The number of rrn ASVs generated by our workflow was comparable to those of ZYMO and PacBio references (**Figure 6B**), indicating that UMI-based error correction did not inflate ASV richness. When benchmarked against the PacBio ASV reference, 84 rrn ASVs were error-free (**Figure 6C**). This level of accuracy is critical for eliminating the “noise” (i.e., sequencing errors) in the ∼4500 bp 16S–ITS–23S rRNA operon profiling, ensuring that variation detected among closely related taxa represents biological divergence rather than technical false signals. In addition, no significant PCR biases associated with GC content in the CCS, forward, and reverse UMI were detected (**Figure S10**).

**Figure 6.**
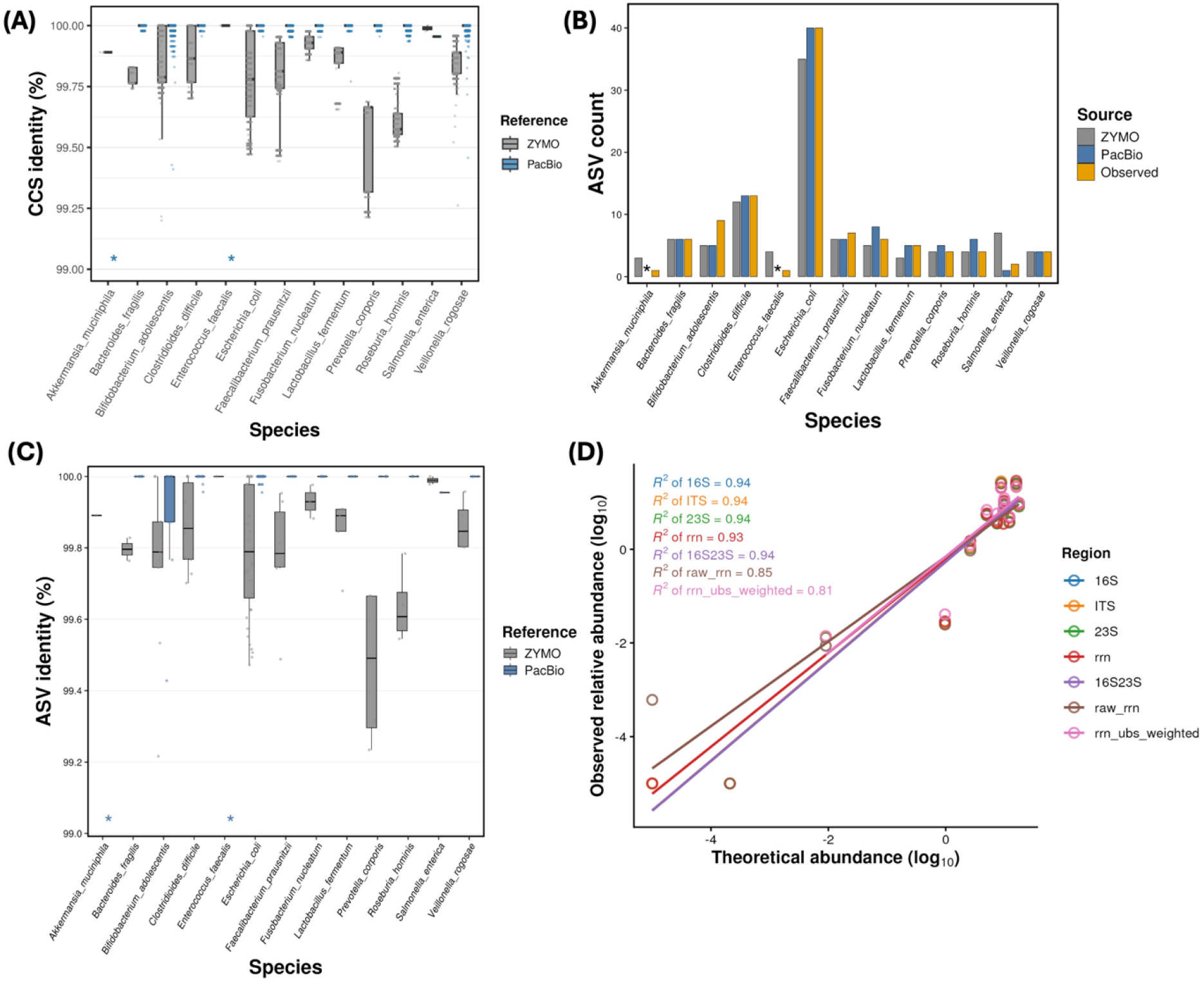
Performance evaluation of UMI-based 16S–ITS–23S rRNA (rrn) consensus sequences (CCS) and amplicon sequence variant (ASV) from the ZymoBIOMICS Gut Microbiome Standard. (A) Distribution of rrn CCS identity to ZYMO (grey) and PacBio (blue) ASV references across species. (B) Species-level rrn ASV counts recovered from the ZYMO reference, PacBio reference, and observed Nanopore sequencing dataset. (C) Distribution of ASV identity to ZYMO (grey) and PacBio (blue) ASV references across species. (D) Observed relative abundance profiles generated from consensus reads using different regions compared with the theoretical composition. Asterisks indicate species for which there is no PacBio-derived ASV reference hit. The 16S, ITS, 23S, rrn, 16S23S, raw_rrn, and rrn_ubs_weighted represent consensus sequences (CCS) of 16S rRNA gene, the internal transcribed spacer, 23S rRNA gene, concatenated 16S+23S rRNA genes, 16S–ITS–23S rRNA operon, the raw reads of rrn, and ccs of rrn weighted by its UMI bin size, respectively.

To evaluate the performance of different markers on detection and quantification, we profiled the mock community composition using 16S, 23S, ITS, 16S23S and rrn. At the species level, the community structures profiled using the CCS of the 16S, ITS, 23S, 16S23S, and rrn were highly correlated with the theoretical composition (R^2^ ≥ 0.93; **Table S5**). In comparison, raw rrn reads yielded lower agreement (R^2^ = 0.85), and polishing alone did not improve quantitative accuracy (rrn_ubs_weighted, R^2^ = 0.81), due to the variable amplification depth (i.e., UMI bin size, ubs) caused by PCR biases, which were also observed in a previous study using amplicon or metagenomic sequencing (Peng and Dorman, 2023; Pu et al., 2025; Smith et al., 2017). Quantification based on CCS yields the highest accuracy because UMI-guided binning collapses PCR duplicates into a single CCS that reflects true molecule counts rather than biased amplification copies. The result demonstrates that the UMI-based error correction and binning workflow contributed to both improved read accuracy and reduced PCR-induced quantitative biases, thereby correcting quantitative distortions and yielding a highly accurate representation of microbial community structure.

In addition to quantitative results, the 16S23S-derived phylogeny, particularly for the five distinct *Escherichia coli* strains, exhibited the highest capability to reconstruct high-resolution phylogenies and recapitulate genome-level evolutionary patterns compared to the other evaluated markers (**Figure 7**). Although these reference units are described as strains by ZYMO based on GTDB taxonomy, each represents a distinct genome. Thus, this analysis can be interpreted as a genome-level assessment of marker performance.

**Figure 7.**
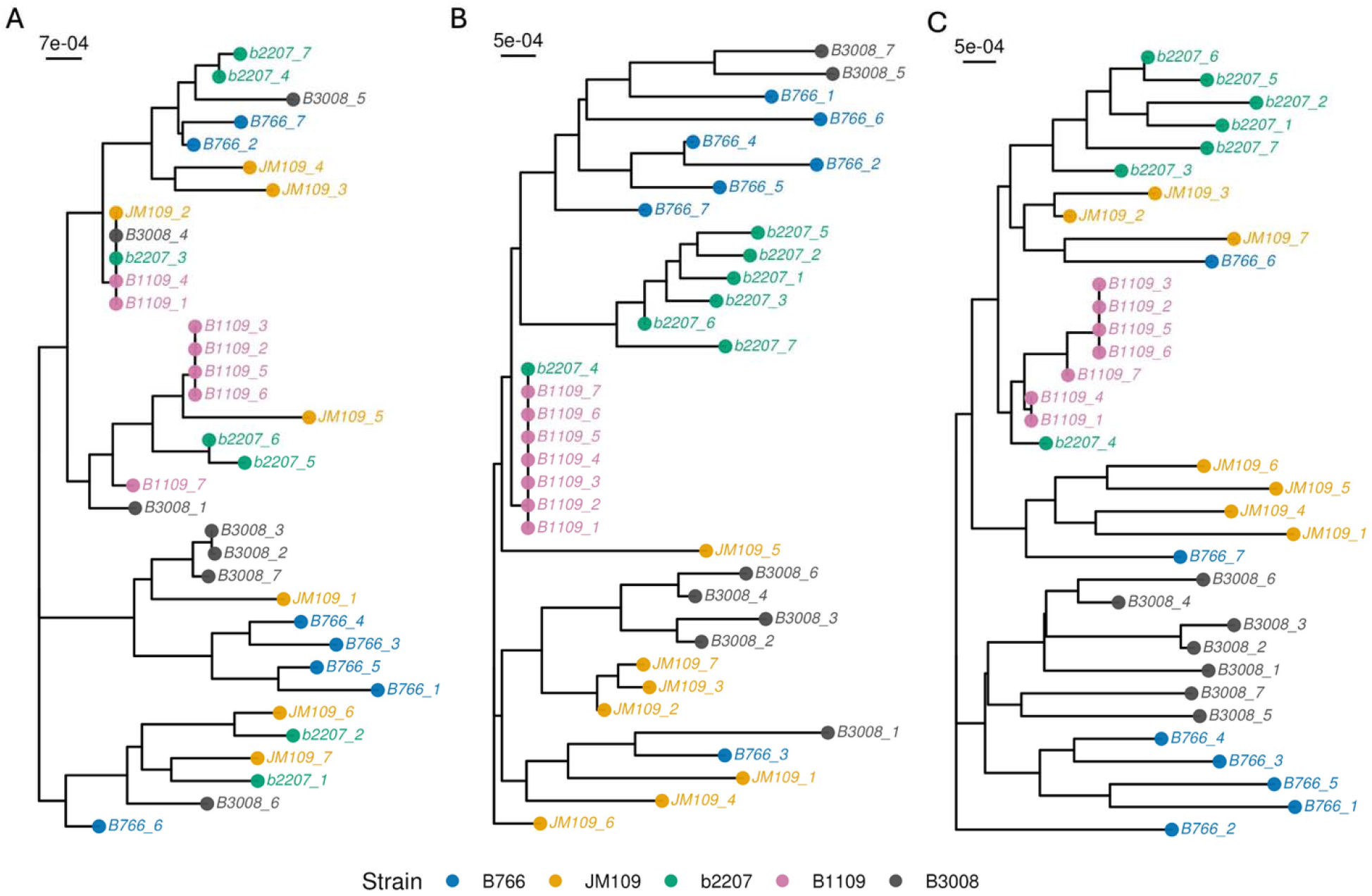
Phylogenetic trees reconstructed from the (A)16S, (B) 23S, and (C) concatenated 16S+23S rRNA genes, extracted from genomes of different *Escherichia coli* strains. Tips represent individual marker copies of each strain and are labeled as strain_copy number. Tip colors indicate different strains across all panels. Tree scale bars are shown in each panel.

Silhouette analysis of patristic distances quantitatively supports the topological observations (**Table S6**) (Rousseeuw, 1987). The 16S23S marker yielded the highest mean silhouette width (0.11), indicating optimal intra-strain sequence identification. Although the 23S marker produced the second-highest mean score (0.03), its intra-genomic variance across individual copies (SD = 0.57) limits its utility as an independent marker for genome-level clustering. The 16S (-0.04), ITS (-0.17), and rrn (-0.15) yielded negative mean silhouette values, suggesting a lack of genome-level resolution. Therefore, from both genome-wide similarity and phylogenetic perspectives, 16S23S is the most informative marker for genome-level inference, followed by 23S rRNA and then 16S rRNA, whereas ITS captures intragenomic heterogeneity rather than genome-level evolutionary signal. The reduced performance of entire *rrn* operon reflects the inclusion of the highly variable ITS region, which inflates topological distance independently of genome-relatedness. Residual inconsistency results from identical copies shared across genomes, which provide no marker-informative variation for phylogenetic or taxonomic separation. For example, identical 23S copies between *E. coli* b2207 (b2207_4) and *E. coli* B1109 (B1109_1∼7) limit the resolution of 23S-based trees (**Figure 7B**). In contrast, the concatenated 16S+23S marker retains additional variation that resolves some of these otherwise indistinguishable copies.

Although the 16S23S marker still cannot fully resolve operons that are completely identical across genomes, such ambiguities could potentially be addressed by incorporating co-abundance-based binning to assign shared marker sequences to their genome (Alneberg et al., 2014).

### 3.4. UltraRes-rrn delivers high-confidence community profiling and ITS-resolved microdiversity in full-scale partial denitrification–anammox reactors

We applied the UltraRes-rrn workflow to evaluate microbial dynamics in two full-scale mainstream PdNA IFAS reactors operating under distinct carbon strategies. Six temporal samples were collected from IFAS 1 (glycerol supplemented with PCE at a total of C/N ratio of 4.29 ± 2.1 g COD/g NO □ □ -N) and four from IFAS 2 (PCE only, testing strict carbon-limited conditions) as reported in a previous study (Bachmann et al., 2025). The DB extraction method successfully yielded a long DNA fragment peak at 17,443 bp (**Figure S11**). Subsequent UMI-based PCR and sequencing generated 61,375–246,730 raw reads and 578–2497 CCS reads per sample (**Table S7**).

RDP classification confidence declined from phylum to species across all samples. At the species level, only the 16S23S and *rrn* markers maintained median confidence scores >0.80 (**Figure 8A**), whereas the median confidence score of ITS decreased under 0.8 at the genus level. Consequently, the 16S23S and *rrn* markers show a median retention rate over 50% at the species level, whereas single short markers (16S, ITS, or 23S) resulted in the rejection of >50% of CCS reads as “unclassified” (i.e., confidence score < 0.8) (**Figure 8B**). Differences in data retention directly impact downstream ecological inferences. While 23S and 16S23S showed comparable species richness, only 16S23S and rrn consistently captured the highest Shannon diversity, accurately reflecting community evenness. Furthermore, PCoA revealed that ITS-derived beta diversity significantly diverged from other markers (**Figure 8C**). Differences in community composition were driven primarily by marker-dependent classification failure rather than conflicting taxonomic assignments. Using 16S23S as the reference marker, we applied the RDP confidence threshold of 0.8 and traced the taxonomic assignment of each sub-region read back to the same parent consensus read. Total discrepancies between short markers, including 16S, ITS, and 23S, and 16S23S were dominated by the 16S23S-only calls category, in which the read was confidently classified by 16S23S markers but fell below the confidence threshold for the comparison marker (**Figure S12**). In contrast, the conflict assignment category, in which the read was classified by both markers but assigned to different species, accounts for only 0-1.6% of total reads. Thus, short-marker profiles differed from 16S23S mainly through read loss after confidence filtering rather than systematic misclassification. The rrn marker is comparable to 16S23S in classification efficacy, but the additional ITS region provided no clear enhancement in taxonomic resolution. Coupled with our earlier findings that the 16S23S marker exhibits a stronger correlation with whole-genome ANI while being significantly less confounded by intragenomic divergence, we identified the 16S23S as the most robust and optimal marker for high-resolution community profiling.

**Figure 8.**
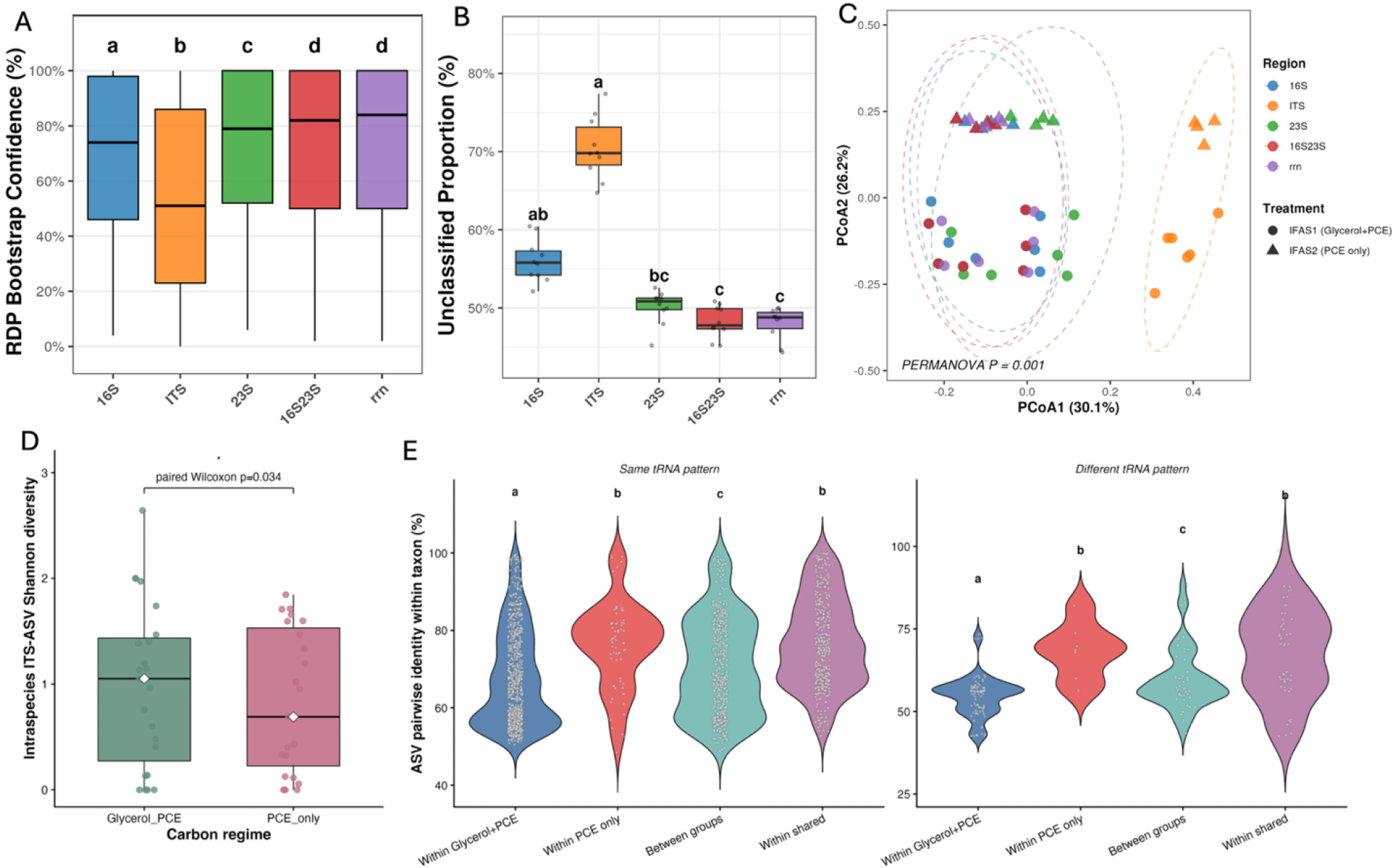
(A) Taxonomic assignment confidence of all consensus sequences and (B) proportion of unclassified sequences among all consensus sequences across different regions (i.e., 16S, ITS, 23S, 16S23S, and rrn). (C) Beta-diversity of microbial community profiled by consensus reads with of RDP bootstrap confidence score over 0.8 of different regions (i.e., 16S, ITS, 23S, 16S23S, and rrn) at the species level. Different lowercase letters indicate statistically significant differences among the genomic regions (*p* < 0.05). Colors indicate the targeted genomic regions (16S, ITS, 23S, 16S23S, and rrn). Point shapes represent the different carbon sources: IFAS 1 (Glycerol+PCE, circles) and IFAS 2 (PCE only, triangles). Dashed ellipses indicate the 95% confidence intervals for each genomic region. The statistical significance of differences in community composition was evaluated using PERMANOVA. (D) Distribution of mean intraspecies ITS ASV Shannon diversity across different carbon regimes. (E) Pairwise sequence identity of ITS ASV sequences within the same species, stratified by same (left) and different (right) tRNA occurrence pattern. Points represent individual ASV pairs, with letters above violins denoting significant differences between categories (P < 0.05). The x-axis classifies pairs into four categories: (1) Within Glycerol+PCE: both ASVs originated from the Glycerol+PCE group; (2) Within PCE only: both ASVs originated from the PCE-only group; (3) Between groups: ASVs in a pair belong to different treatment groups; and (4) Within mixed samples: represents overall within-taxon variation for taxa shared by both groups, regardless of group origin.

In the two IFAS reactors, high-resolution 16S23S profiling revealed clear carbon-source-driven community divergence (**Figure S13**), while alpha diversity remained broadly comparable between reactor types (**Figure S14**). The pattern was also observed with the other markers. However, different markers infer different core bacteria among each group, and only 16S23S and rrn are consistent (**Figure S15**). As shown in **Figure S16**, both systems supported the PdNA pathway, with *Brocadia sapporoensis* identified as a core anammox (AMX) species by all markers (Meng et al., 2022; Qiao et al., 2025; Zhang et al., 2025). However, the markers differed in their ability to resolve between-system variation. Specifically, markers other than ITS showed that the Glycerol+PCE reactor maintained a higher relative abundance of *B. sapporoensis* and also harbored a secondary AMX species, *Brocadia pituitae*, which was rare in the PCE-only reactor, consistent with carbon limitation (Zhao et al., 2026). By contrast, ITS alone failed to detect *B. pituitae*, indicating incomplete species-level recovery and reduced sensitivity to ecologically relevant differences between systems. Although ITS alone has limited ecological clarity, integrating the ITS as a secondary marker with its corresponding 16S23S-defined species or genome enables its use as a secondary marker for fine-scale ecological inference for microdiversity and intragenomic diversity (Brown and Fuhrman, 2005).

At the same-species level, mean ITS ASV Shannon diversity at intraspecies level was higher under the Glycerol+PCE regime (**Figure 8D**). Consistently, ITS ASV pairwise identity stratified by ITS tRNA pattern showed greater divergence among taxa unique to the Glycerol+PCE reactor (**Figure 8E**). This supports the environmental filtering and the coexistence of closely related ecotypes under carbon-replete conditions (Larkin and Martiny, 2017; Stewart and Cavanaugh, 2007a). These findings highlight the value of coupling high-accuracy full-length rRNA consensus reads with a hierarchical marker strategy. The conserved 16S23S marker resolved microbial composition shift driven by carbon-regime, whereas the hypervariable ITS captured within-taxon microdiversity that may reflect niche adaptation. Overall, this framework revealed PdNA community assembly patterns that would be missed by conventional 16S rRNA gene amplicon sequencing. By linking carbon availability to both community-level turnover and ecotype-level variation, UltraRes-rrn provides a high-resolution view of how operational conditions shape microbial diversity and reactor stability.

## Conclusion

In this study, we propose the hierarchical utilization of the 16S–ITS–23S rRNA operon as the marker and developed the UltraRes-rrn framework, offering a robust solution for high-resolution microbial profiling in complex environmental samples. By integrating HMW DNA extraction, UMI-based Nanopore sequencing, and a tailored analytical pipeline, we bridged the gap between low-accuracy long reads and high-resolution taxonomic requirements. Our approach demonstrated the feasibility of capturing not only species-level diversity but also microdiversity and intragenomic variations that are challenging to traditional amplicon gene sequencing. In addition, the UMI-based method enables the mitigation of PCR-biases, supporting high accuracy quantitation. Overall, this framework provides a cost-effective, high-accuracy alternative to metagenomic shotgun sequencing for researchers primarily focused on high-resolution community composition. Despite these improvements, several challenges remain to be addressed in the future. First, while we increased genomovar-level resolution via ASV-resolved rRNA operons, confidently partitioning the phylogenetic tree to cluster multiple rrn copies within a single genome remains complex. Second, while our amplicon-based approach is efficient, broader validation across diverse sample types is necessary. Integrating more whole-genome metagenomic sequencing data will be crucial to benchmarking our results and clarifying the ecological representativeness of our workflow.

## Notes

### Competing Interest Statement

The authors have declared no competing interest.

