## Supplemental information for "High-accuracy hierarchical rRNA operon profiling resolves genomovar-level taxonomy and microdiversity using Nanopore sequencing"

### Text:

Text S1. Summary of tRNA gene occurrence pattern.

Across all genomes, a total of 286,017 tRNA genes were detected, spanning 20 tRNA types. The distribution was dominated by tRNA-Ala (118,066) and tRNA-Ile (106,602), followed by tRNA-Glu (55,614). Other types were less frequent (e.g., tRNA-Val: 2,732, tRNA-Lys: 2,630).

### Figures:

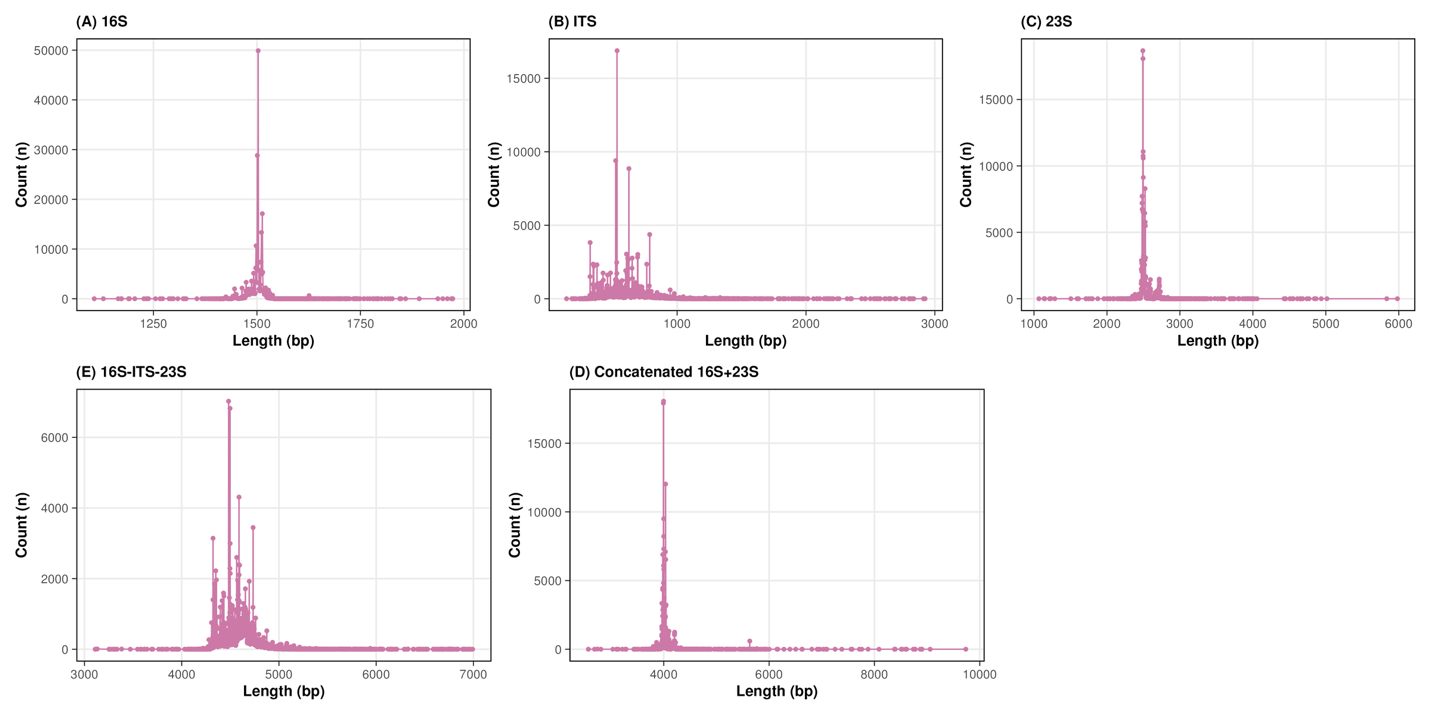

Figure S1. Length distribution of (A) 16S, (B) ITS, (C) 23S, (D) 16S-ITS-23S rRNA operon, and (E) concatenated 16S+23S rRNA gene in the rRNA operon datasets generated.

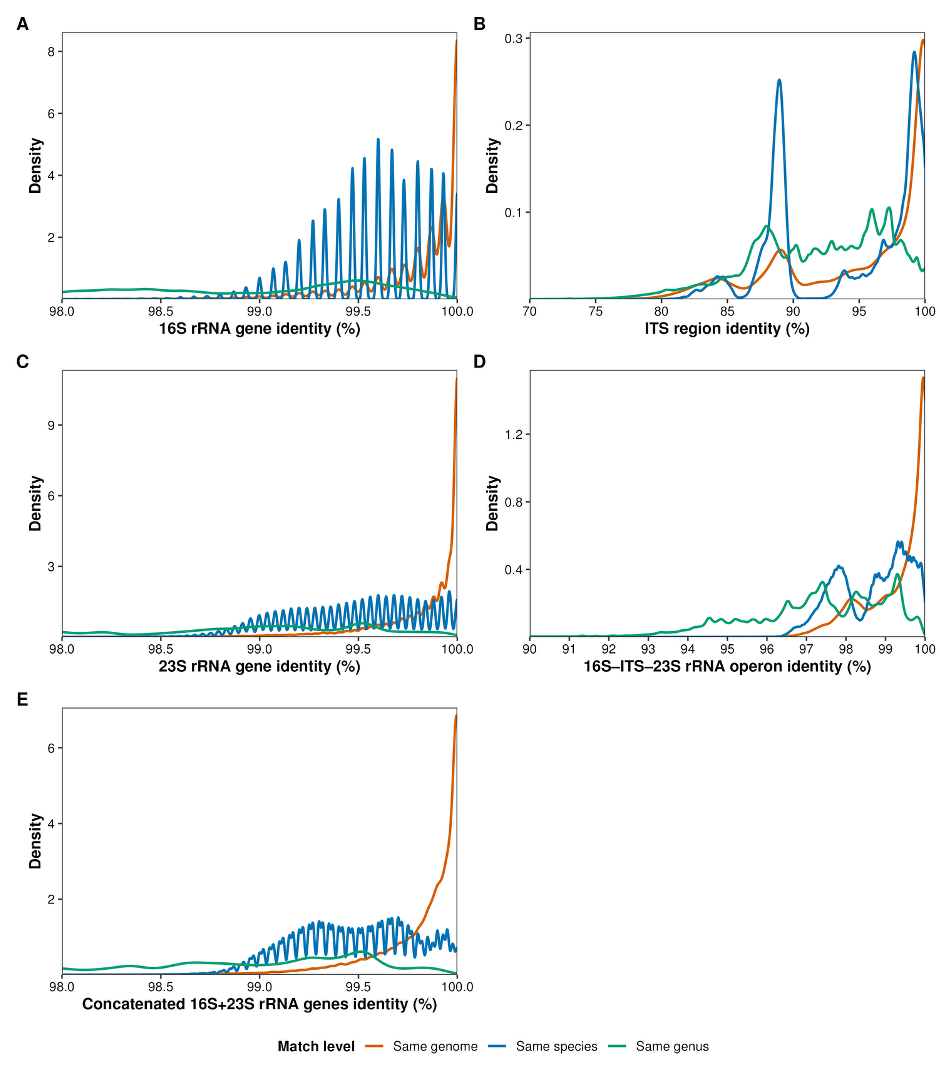

Figure S2. Pairwise identity of (A) 16S, (B) ITS, (C) 23S, (D) concatenated 16S+23S rRNA gene, and (E) 16S-ITS-23S rRNA operon within the same genome, species, and genus, which are defined by GTDB taxonomy.

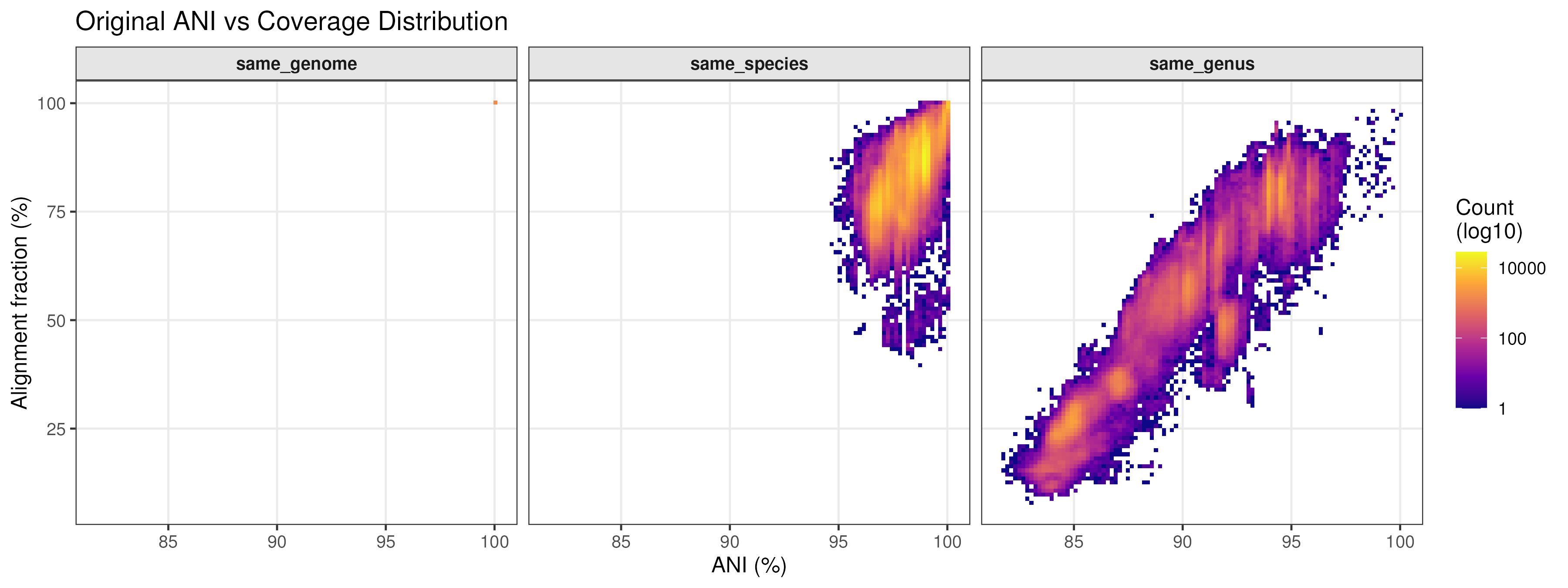

Figure S3. Alignment Fraction (AF) and Average Nucleotide Identity (ANI) distributions for genome pairs classified within the same strain, species, and genus, as defined by the Genome Taxonomy Database (GTDB).

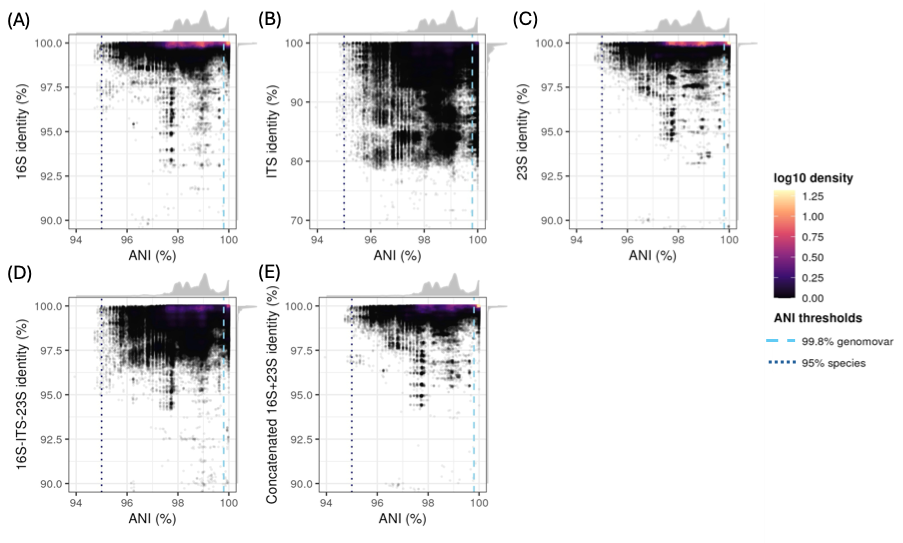

Figure S4. Full-scale density plot of correlation between whole-genome Average Nucleotide Identity (ANI) and pairwise identity of (A) 16S, (B) ITS, (C) 23S, (D) concatenated 16S+23S rRNA gene, and (E) 16S-ITS-23S rRNA operon within the same species, which defined by GTDB taxonomy. The color gradient represents the density of data points at each position. Vertical dash indicates the ANI thresholds of 95%and 99.8% for species and genomovar level designations, respectively.

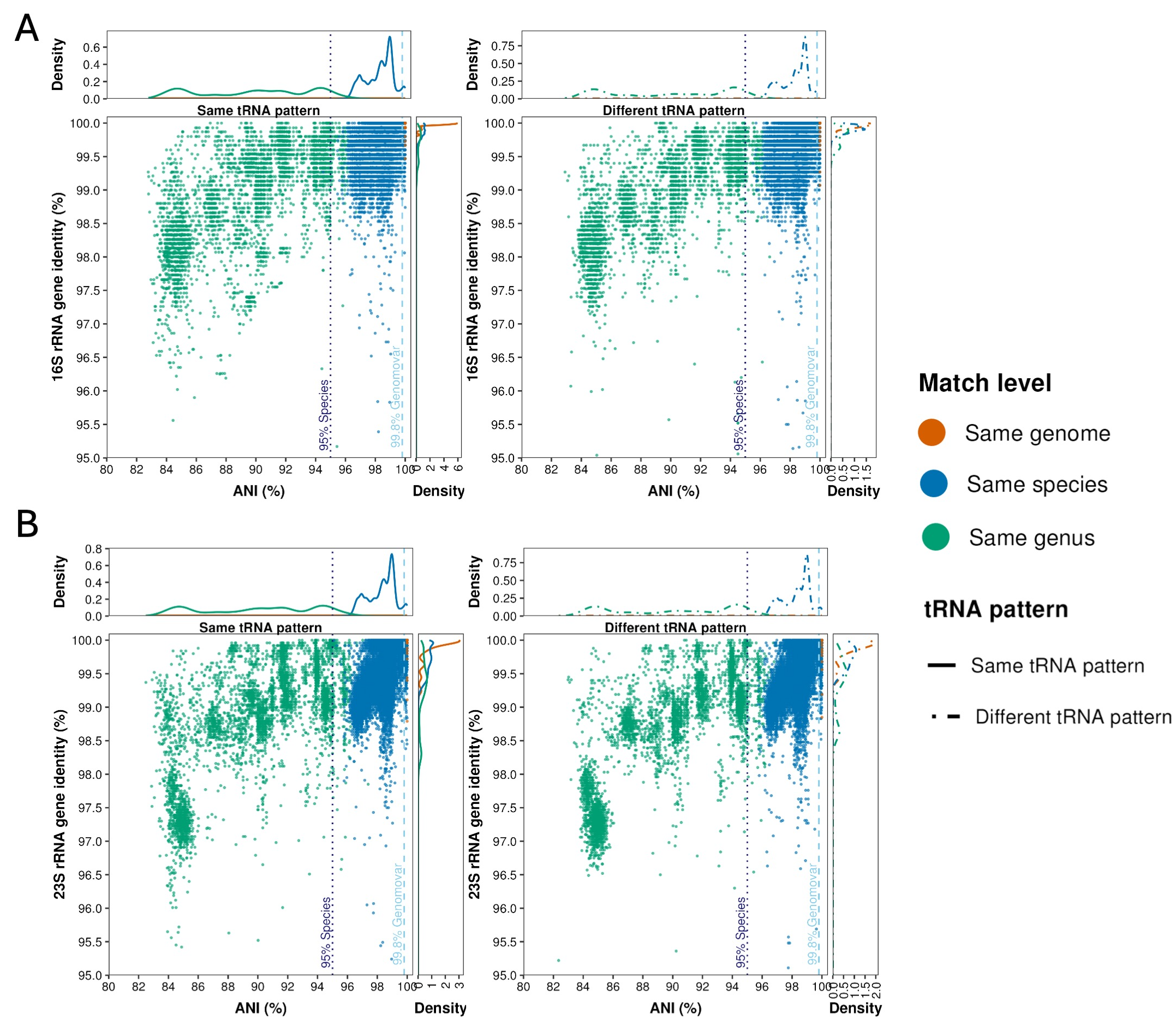

Figure S5. Density plots of whole-genome ANI versus pairwise sequence identity across rRNA regions, stratified by tRNA gene occurrence pattern concordance for (A) 16S and (B) 23S rRNA genes. In each panel, the left and right plots represent the sequence comparisons with the same or different tRNA occurrence patterns, respectively. The top and right marginal density curves show the univariate distribution of ANI and region-specific identity, respectively. Point colors in the main scatterplots and line colors in the marginal density plots indicate the taxonomic relationship of each sequence comparison, as defined by GTDB taxonomy. Vertical dashed lines indicate whole-genome ANI thresholds of 95.0% and 99.8%, corresponding to commonly used species- and genomovar-level genomic boundaries, respectively.

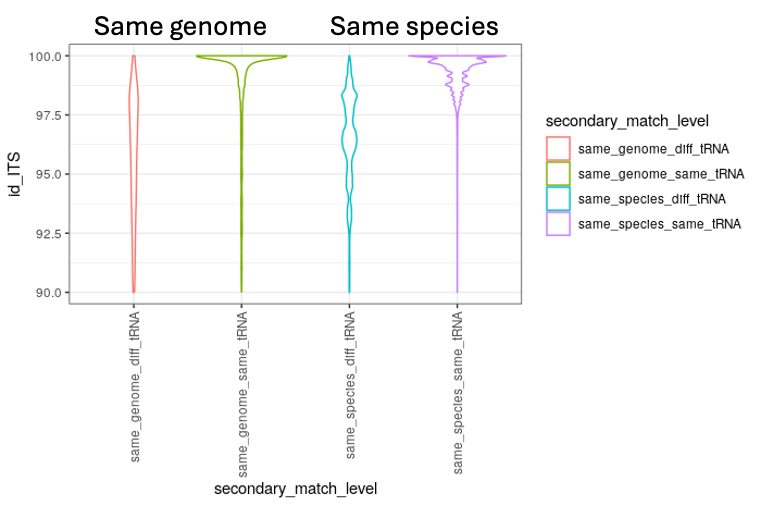

Figure S6. Distribution of identity of ITS with different or the same tRNA patterns within the same genomes or species.

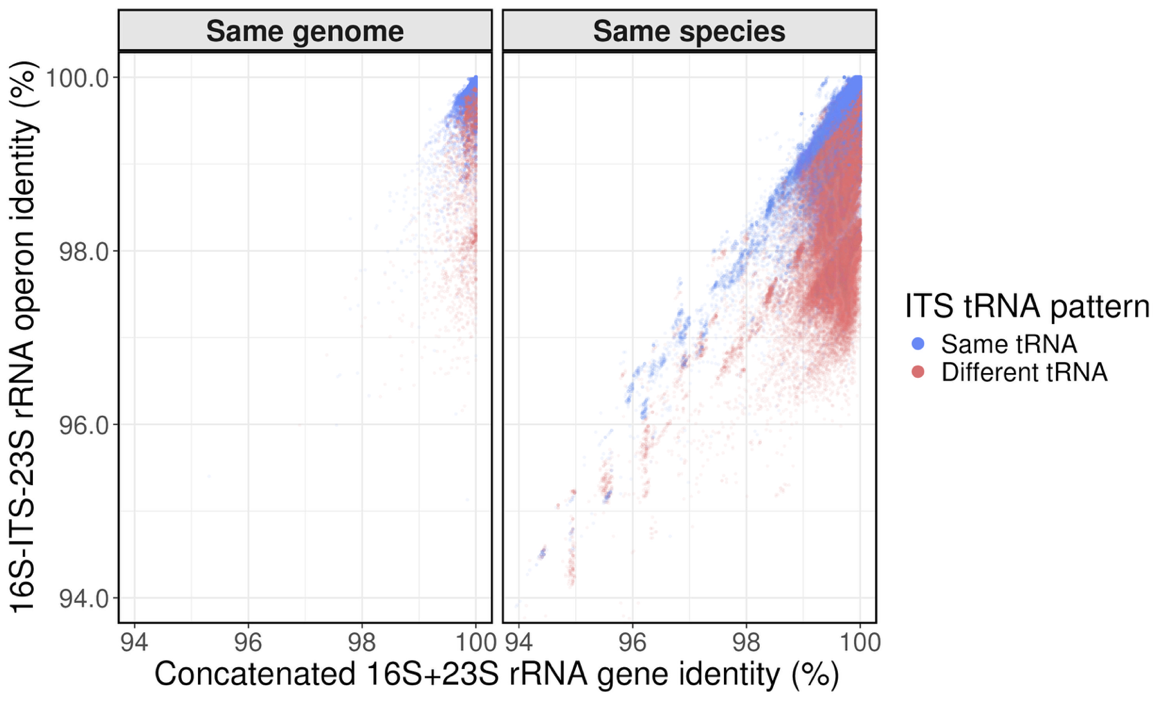

Figure S7. Comparison of the pairwise identity between the 16S-ITS-23S rRNA operon and concatenated 16S+23S rRNA genes at same genome and same species (markers from same species but not same genome) levels. Colors indicate whether each sequence pair shares the same or different tRNA occurrence patterns.

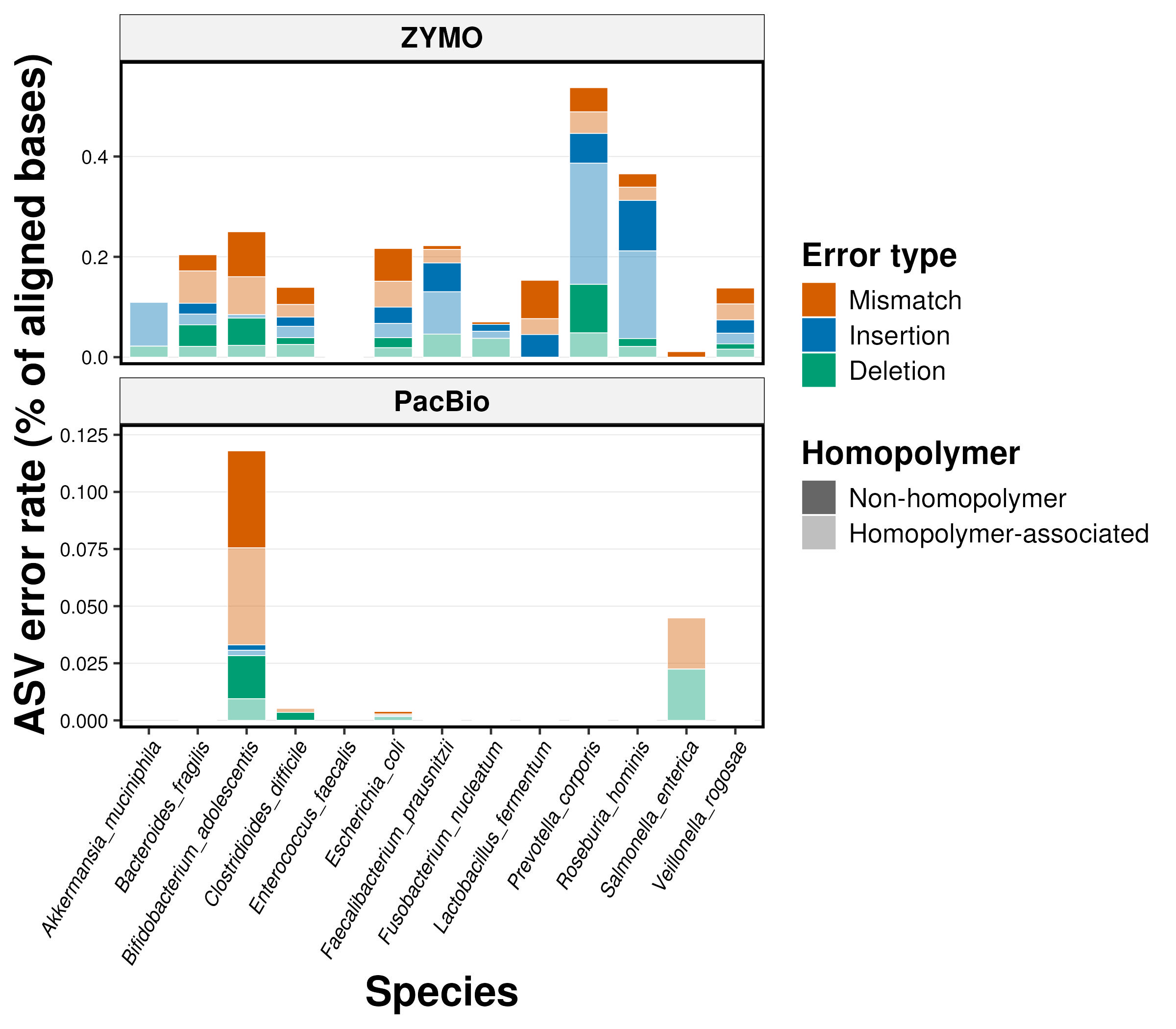

Figure S8. Error profiles of consensus reads against ZYMO (top panel) and PacBio (bottom panel) reference databases. Colors represent different error types, including mismatch (orange), insertion (blue), and deletion (green). Light and dark shades indicate homopolymer- and non-homopolymer-associated errors, respectively.

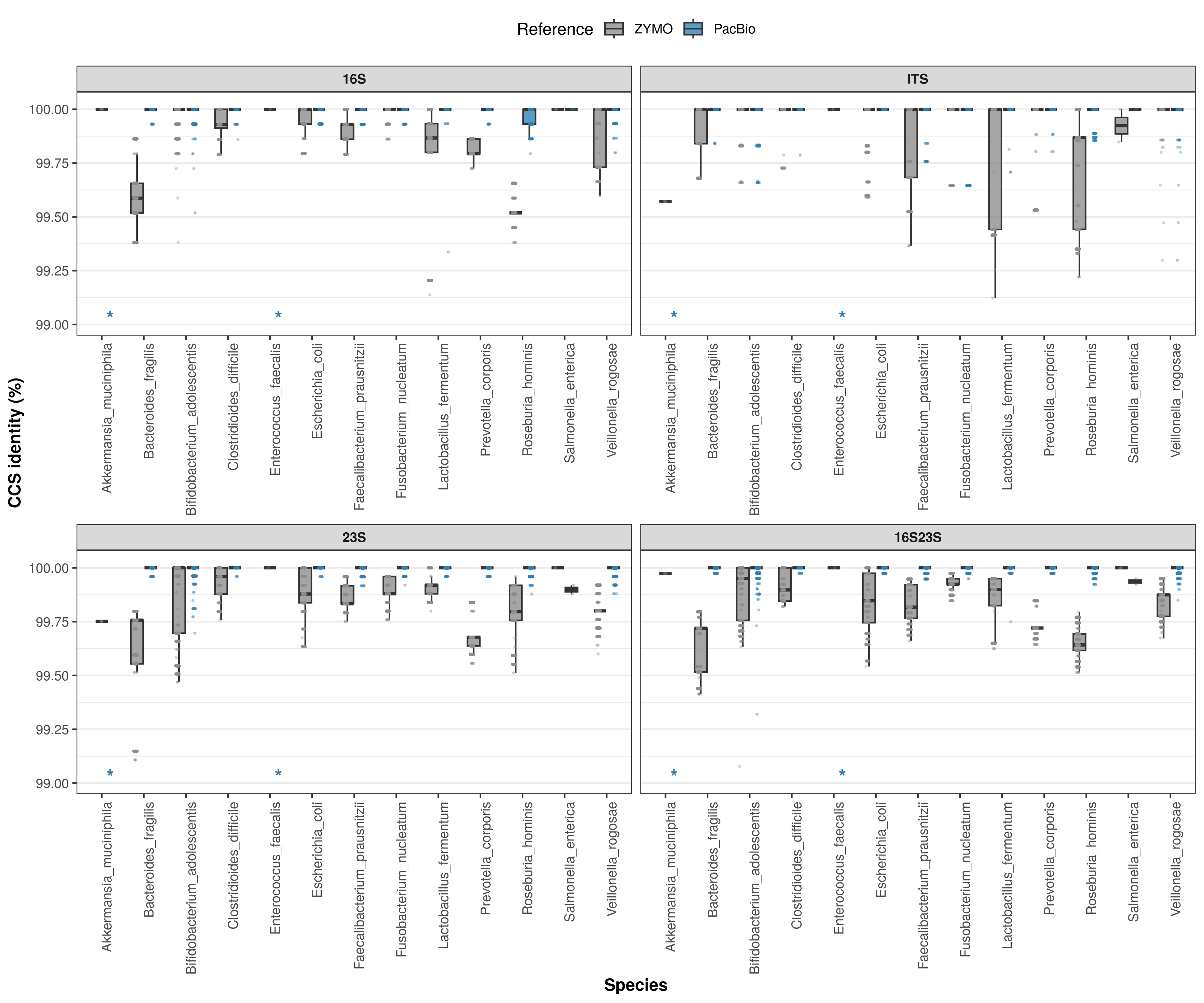

Figure S9. Distribution of CCS identity to ZYMO and PacBio ASV references across species for 16S, ITS, 23S, and 16S23S regions. Colors of boxplots represent the ZYMO- (grey) and PacBio-derived (blue) reference database. Asterisks indicate species for which there is no PacBio-derived ASV reference hit.

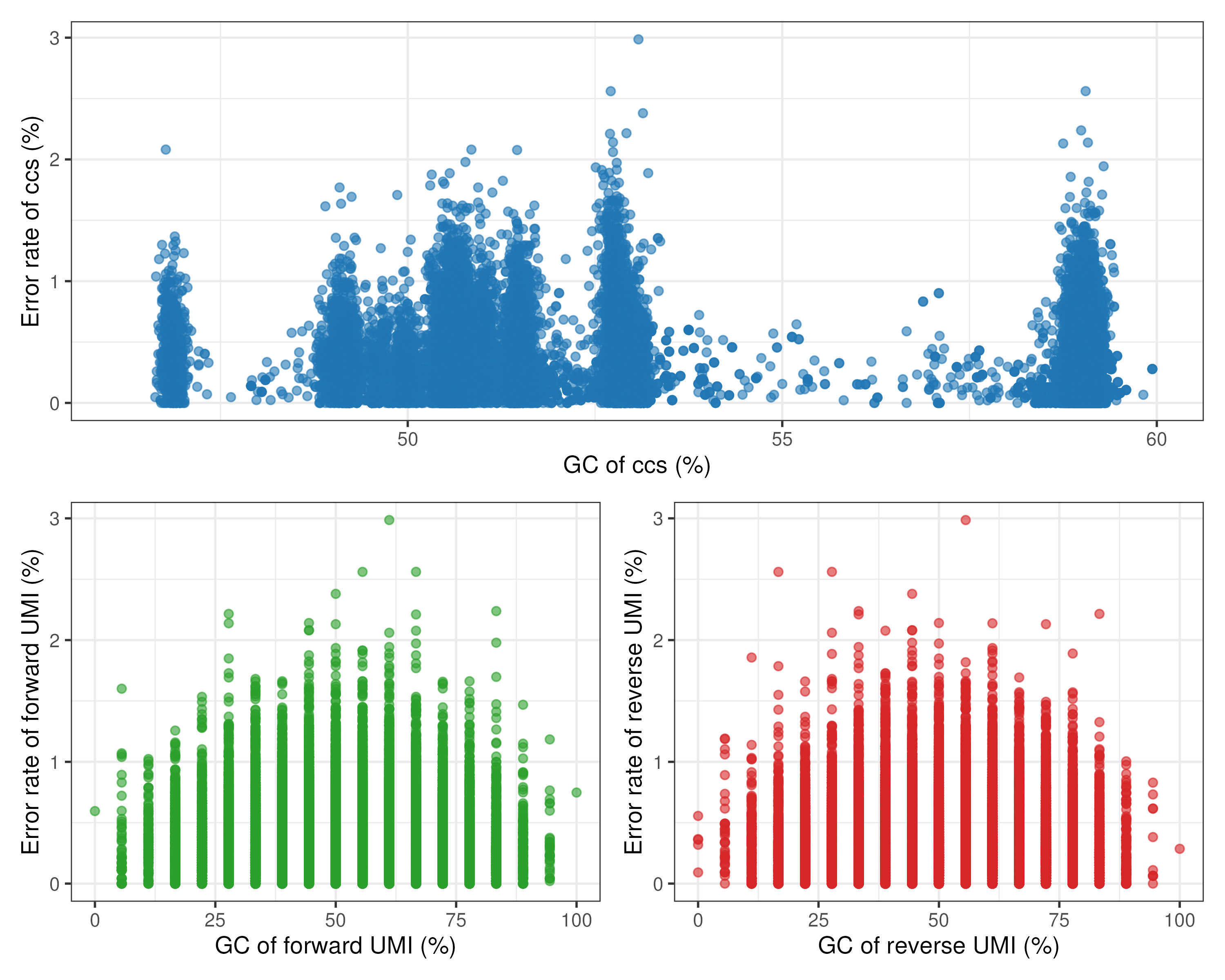

Figure S10. Error rate distribution across varying GC contents for consensus sequences (CCS), forward and reverse unique molecular identifiers (UMI).

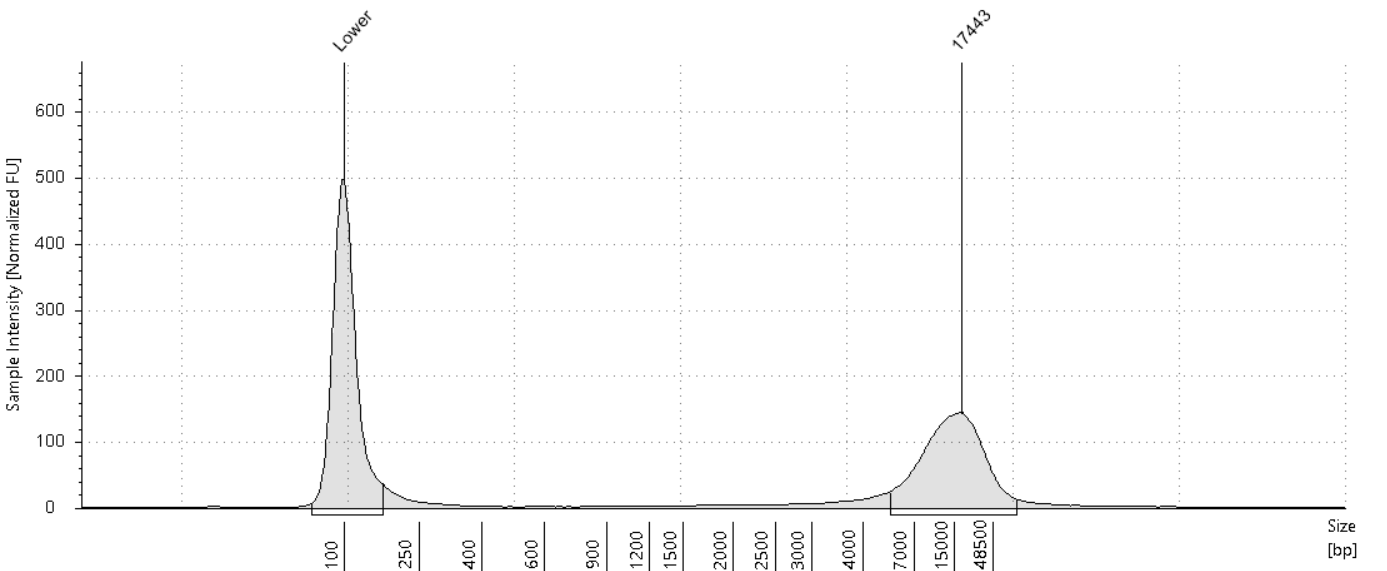

Figure S11. Fragment distribution of environmental samples. Peaks annotated with Lower indicate the Lower Marker in the TapeStation analysis.

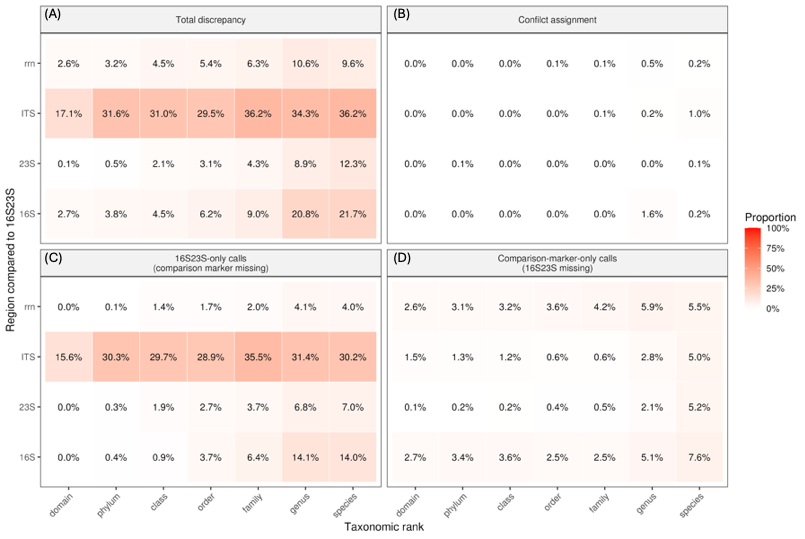

Figure S12. Pairwise discrepancy heatmaps comparing taxonomic assignments at read level from different marker regions against the 16S23S as a reference. Heatmaps summarize region-wise disagreement patterns relative to 16S23S across taxonomic ranks, with consensus reads from all samples combined. Rows indicate the comparison region (16S, ITS, 23S, and rrn), and columns indicate taxonomic rank. Four discrepancy categories indicate the proportion of consensus sequence in which (A) total discrepancy rate, including both conflict assignment and missing classification. (B) 16S23S and the comparison region were both classified, but the annotation taxon is different (True discordance among all reads). (C) 16S23S was classified but the comparison marker was missing (16S23S called, region missing). (D) The comparison marker was classified but 16S23S was missing (region called, 16S23S missing). For each region–rank comparison, individual reads were first assigned to one of these read-level comparison outcomes, and cell values represent the percentage of total reads obtained by summing read counts within the corresponding category. Color intensity reflects the same proportion. Taxonomic classifications were compared using calls passing the specified RDP confidence threshold.

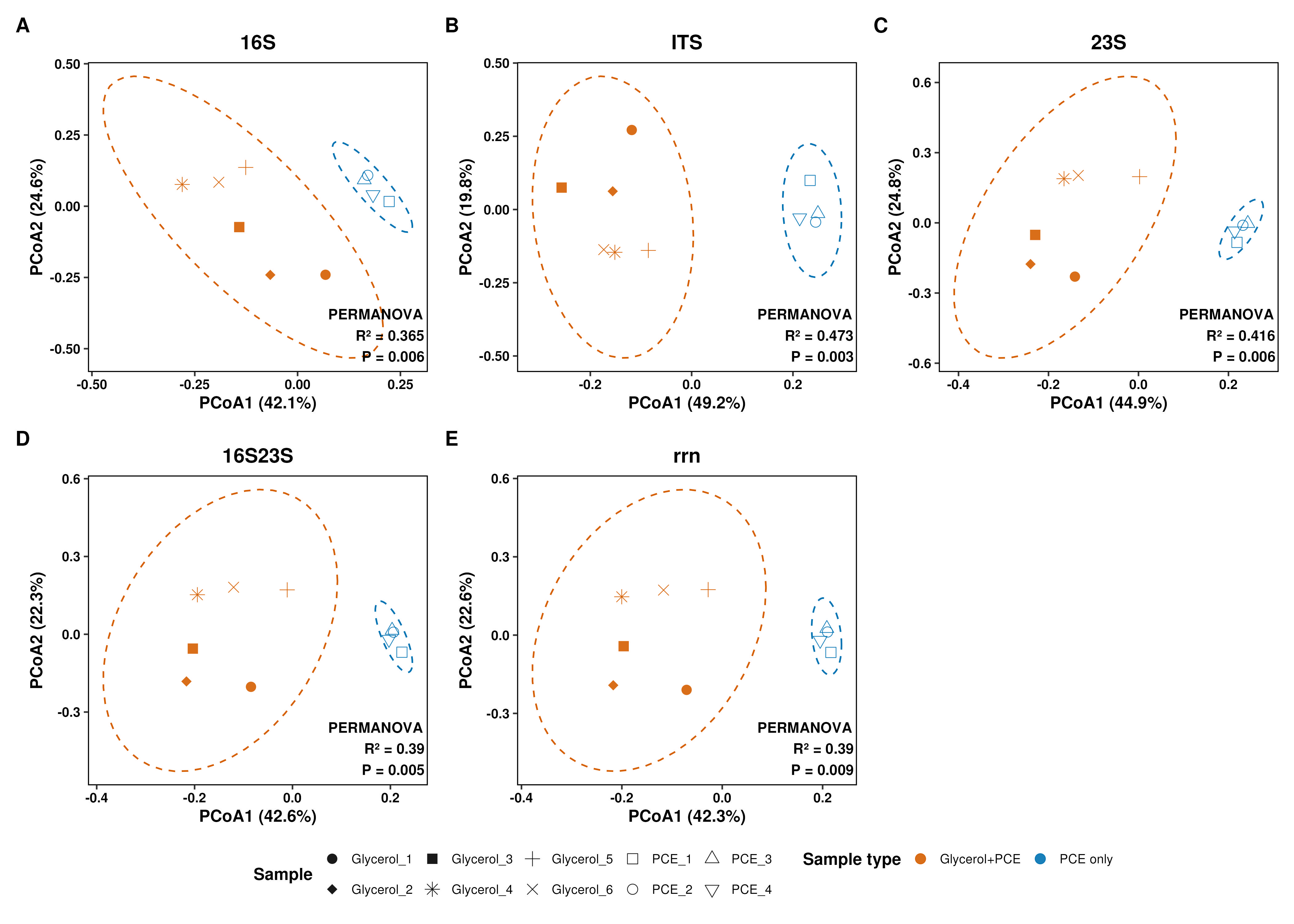

Figure S13. Beta diversity of microbial communities across different markers (A) 16S, (B) ITS, (C) 23S, (D) 16S23S, and (E) rrn regions. Principal Coordinate Analysis (PCoA) plots based on Bray-Curtis distances illustrating the structural variations of microbial communities of different type of samples with a confidence score cutoff of 0.8. Colors indicate different type of samples. Point shapes represent the different samples. Dashed ellipses indicate the 95% confidence intervals for each genomic region. The statistical significance of differences in community composition was evaluated using PERMANOVA, with *p*-values provided in each panel.

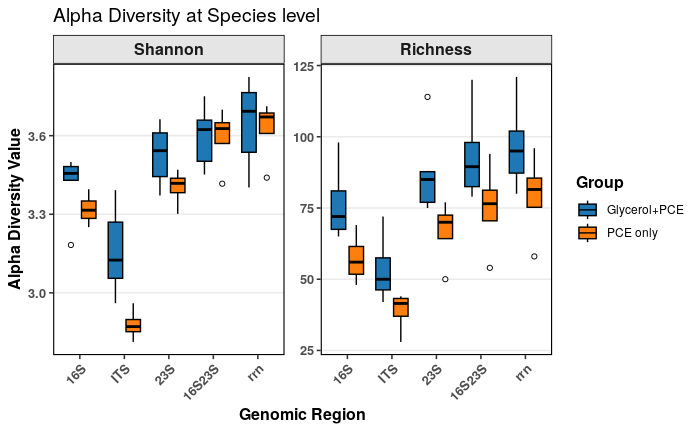

Figure S14. Alpha diversity of microbial communities of different types of samples across different markers. Left and right panels indicate the Shannon and Richness diversity, respectively. Colors of the boxplot represent different types of samples. 16S, ITS, 23S, 16S23S, and rrn on the X-axis indicate the different markers.

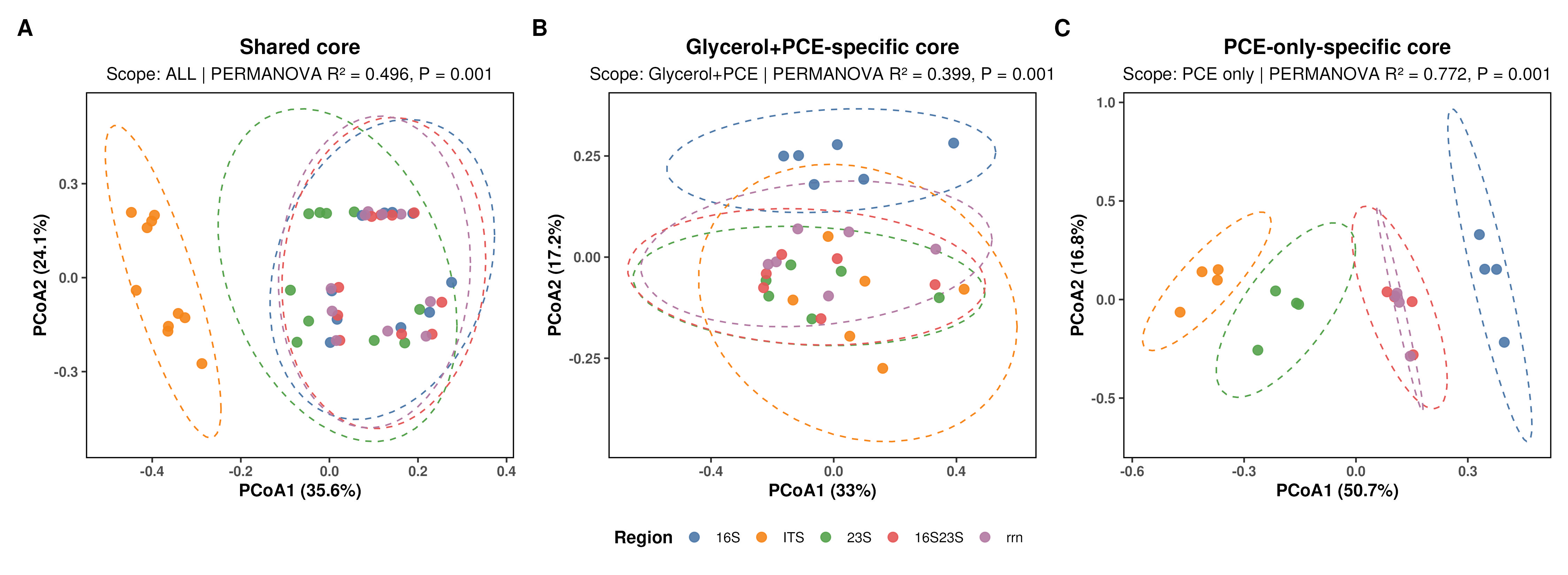

Figure S15. Beta diversity of core microbial communities (A) shared between type of samples, (B) IFAS1 (Glycerol+PCE), and (C) IFAS2 (PCE-only) across different markers. Principal Coordinate Analysis (PCoA) plots based on Bray-Curtis distances illustrating the structural variations of microbial communities at different taxonomic levels with a confidence score cutoff of 0.8. Colors indicate different markers (16S, ITS, 23S, 16S23S, and rrn). Dashed ellipses indicate the 95% confidence intervals for each genomic region. The statistical significance of differences in community composition was evaluated using PERMANOVA, with *p*-values provided in each panel.

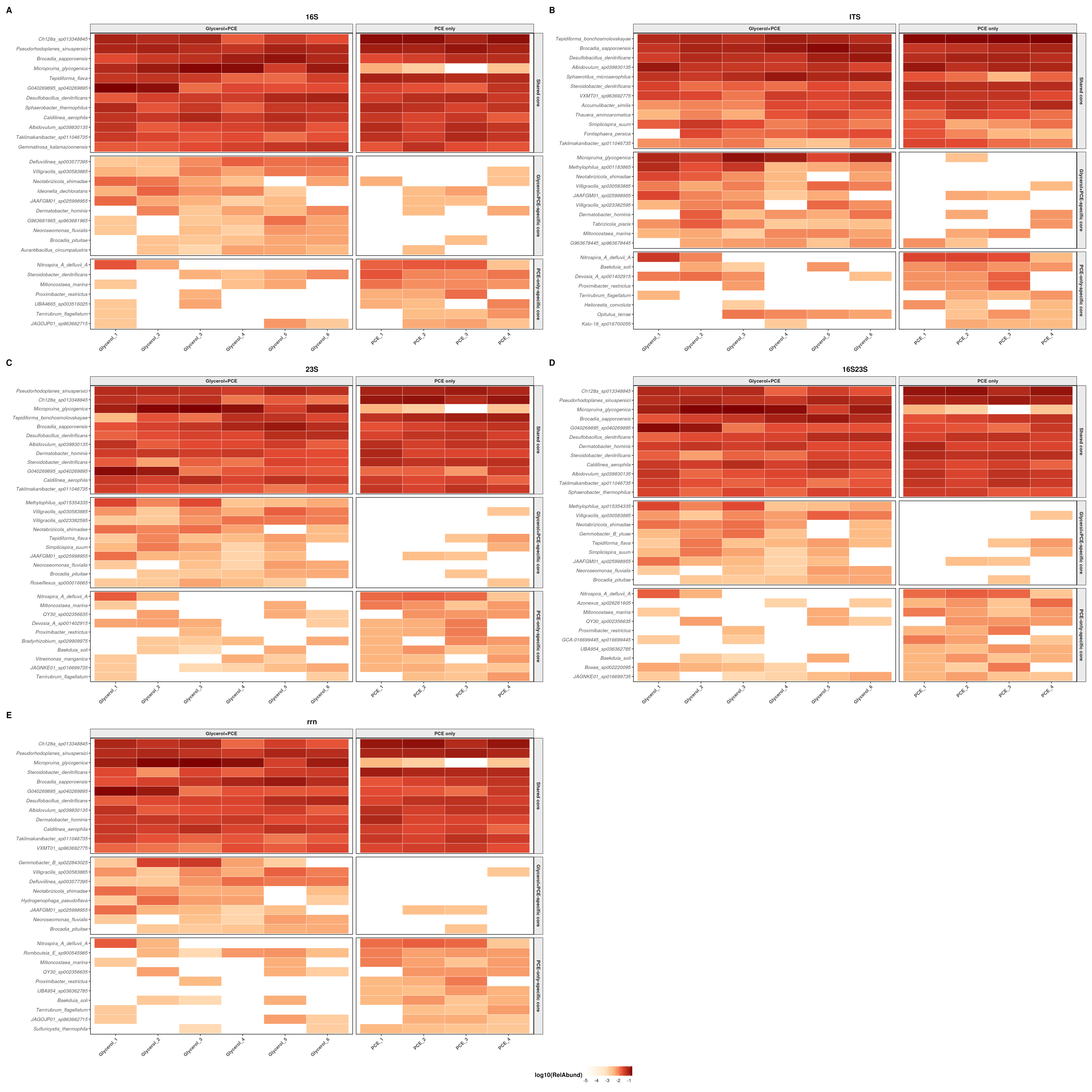

Figure S16. Core species of microbial community profiled by consensus sequences of the (A) 16S, (B) ITS, (C) 23S, (D) 16S23S, and (E) rrn regions with of RDP bootstrap confidence score over 0.8 for samples within different sample types. Sample types (Glycerol+PCE and PCE only) represent the varying carbon regimes in the reactors. Colors indicate the log10-transformed relative abundance of the identified taxa. In each panel, top, middle, and bottom plots indicate the core species shared by two sample types, solely identified in Glycerol+PCE, and solely identified in PCE-only, respectively.

### Table:

Table S1. Sequences of primers used for in silico PCR analysis.

| **Target region** | **Name** | | **Primer sequence 5' to 3'** | **Min length (bp)** | **Max length (bp)** |
| --- | --- | --- | --- | --- | --- |
| 16S rRNA gene | forward | | AGRRTTYGATYHTDGYTYAG | 1000 | 2000 |
|  | reverse | | TACCTTGTTACGACTT |  |  |
| ITS rRNA gene | forward_1 | AAGTCGTAACAAGGTARCY | | 0 | 2000 |
|  | forward_2 | AAGTCGTAACAAGGTA | |  |  |
|  | revserse_1 | CYGAATGGGGVAACC | |  |  |
|  | revserse_2 | CCGAATGAGGAAAAT | |  |  |
| 23S rRNA gene | forward | | GGTTBCCCCATTCRG | 1000 | 6000 |
|  | reverse_1 | | CCRAMCTGTCTCACGACG |  |  |
|  | revserse_2 | | CGTCGTGAGACAGTTCGG |  |  |
| 16S-ITS-23S rRNA operon | forward | | AGRRTTYGATYHTDGYTYAG | 3000 | 7000 |
|  | reverse | | CCRAMCTGTCTCACGACG |  |  |

Table S2. Sequences of primers used in the full rRNA operon unique molecular identifier (UMI) tagging and PCR amplification (Karst et al., 2021).

| **Step** | **Name** | **Primer sequence 5' to 3'** |
| --- | --- | --- |
| UMI tagging | UMI_16S_27F | CAAGCAGAAGACGGCATACGAGATNNNYRNNNYRNNNYRNNNAGRRTTYGATYHTDGYTYAG |
|  | UMI_23S_U2428R | AATGATACGGCGACCACCGAGATCNNNYRNNNYRNNNYRNNNCCRAMCTGTCTCACGACG |
| PCR amplification | UMI_PCR_fw | CAAGCAGAAGACGGCATACGAGAT |
|  | UMI_PCR_rv | AATGATACGGCGACCACCGAGATC |

Table S3. Thermocycling conditions for UMI-tagging and PCR amplification.

| **Step** | **Substep** | **Time** | **Temperature** | **Cycles** | **Reaction volume** |
| --- | --- | --- | --- | --- | --- |
| **UMI tagging** | Initial denaturation | 3 min | 94°C | 1 | 20 µL |
|  | Denaturation | 30 s | 94°C | 2 |  |
|  | Annealing | 15 s | 60°C |  |  |
|  | Extension | 3 min | 68°C |  |  |
|  | Final extension | 10 min | 68°C | 1 |  |
| **PCR-1** | Initial denaturation | 3 min | 94°C | 1 | 20 µL |
|  | Denaturation | 15 s | 94°C | 25 |  |
|  | Annealing | 15 s | 60°C |  |  |
|  | Extension | 3 min | 68°C |  |  |
|  | Final extension | 10 min | 68°C | 1 |  |
| **PCR-2** | Initial denaturation | 3 min | 94°C | 1 | 20 µL |
|  | Denaturation | 15 s | 94°C | 5 |  |
|  | Annealing | 15 s | 60°C |  |  |
|  | Extension | 3 min | 68°C |  |  |
|  | Final extension | 10 min | 68°C | 1 |  |

Table S4. Summary of plastic biofilm carrier sampling from two full-scale Integrated Fixed-film Activated Sludge (IFAS) partial denitrification-anammox (PdNA) reactors (IFAS1 and IFAS2) at the James River Treatment Plant (JRTP).

| **Sample ID** | | **Date Sampled** |  | **Sampling Date** | **Carbon Source** |
| --- | --- | --- | --- | --- | --- |
| **IFAS1 Primary Clarifier Effluent (PCE) + Glycerol** | | | | | |
| **G1** | **Glycerol_1** | 2/27/23 |  | day 294 from startup | Glycerol and PCE carbon |
| **G2** | **Glycerol_2** | 3/28/23 |  | day 323 from startup | Glycerol and PCE carbon |
| **G3** | **Glycerol_3** | 5/5/23 |  | day 361 from startup | Glycerol and PCE carbon |
| **G4** | **Glycerol_4** | 10/17/23 |  | day 526 from startup | Glycerol and PCE carbon |
| **G5** | **Glycerol_5** | 11/20/23 |  | day 560 from startup | Glycerol and PCE carbon |
| **G6** | **Glycerol_6** | 12/28/23 |  | day 598 from startup | Glycerol and PCE carbon |
| **IFAS2 Primary Clarifier Effluent (PCE)** | | | | | |
| **P1** | **PCE_1** | 10/17/23 |  | day 169 from startup | PCE carbon |
| **P2** | **PCE_2** | 11/9/23 |  | day 192 from startup | PCE carbon |
| **P3** | **PCE_3** | 11/20/23 |  | day 203 from startup | PCE carbon |
| **P4** | **PCE_4** | 12/28/23 |  | day 241 from startup | PCE carbon |

Table S5. Composition of ZymoBIOMICS Gut Microbiome Standard.

| Abbreviation | Taxa | Theoretical Composition (%) | | | |
| --- | --- | --- | --- | --- | --- |
|  |  | Genomic  DNA | 16S Only | Genome Copy | Cell Number |
| AM | *Akkermansia muciniphila* | 1.5 | 0.97 | 1.62 | 1.62 |
| BF | *Bacteroides fragilis* | 14 | 9.94 | 8.33 | 8.36 |
| BA | *Bifidobacterium adolescentis* | 6 | 8.78 | 8.83 | 8.86 |
| CD | *Clostridioides difficile* | 1.5 | 2.62 | 1.1 | 1.1 |
| CP | *Clostridium perfringens* | 0.0001 | 0.0002 | 0.00009 | 0.00009 |
| EF | *Enterococcus faecalis* | 0.001 | 0.0009 | 0.0011 | 0.0011 |
| ECB1109 | *Escherichia coli (B-1109)* | 2.8 | 2.46 | 1.77 | 1.77 |
| ECB2207 | *Escherichia coli (B-2207)* | 2.8 | 2.29 | 1.64 | 1.65 |
| ECB3008 | *Escherichia coli (B-3008)* | 2.8 | 2.53 | 1.82 | 1.82 |
| ECB766 | *Escherichia coli (B-766)* | 2.8 | 2.31 | 1.66 | 1.66 |
| ECJM109 | *Escherichia coli (JM109)* | 2.8 | 2.53 | 1.82 | 1.83 |
| FP | *Faecalibacterium prausnitzii* | 14 | 17.63 | 14.77 | 14.82 |
| FN | *Fusobacterium nucleatum* | 6 | 7.49 | 7.53 | 7.56 |
| LF | *Lactobacillus fermentum* | 6 | 9.63 | 9.68 | 9.71 |
| MS | *Methanobrevibacter smithii* | 0.1 | 0.066 | 0.17 | 0.17 |
| PC | *Prevotella corporis* | 6 | 4.98 | 6.26 | 6.28 |
| RH | *Roseburia hominis* | 14 | 9.89 | 12.43 | 12.47 |
| SE | *Salmonella enterica* | 0.01 | 0.009 | 0.007 | 0.0065 |
| VR | *Veillonella rogosae* | 14 | 15.87 | 19.94 | 20.01 |

Table S6. Summary of silhouette scores evaluating the clustering performance of various ribosomal RNA genomic regions. Metrics include mean, median, standard deviation, minimum, and maximum scores.

| Region | mean_silhouette | median_silhouette | sd_silhouette | min_silhouette | max_silhouette |
| --- | --- | --- | --- | --- | --- |
| 16S rRNA gene | -0.04 | -0.13 | 0.39 | -0.57 | 0.70 |
| Internal transcribed spacer | -0.17 | -0.18 | 0.08 | -0.35 | -0.06 |
| 23S rRNA gene | 0.03 | -0.24 | 0.57 | -1.00 | 1.00 |
| Concatonated 16S+23S rRNA genes | 0.11 | -0.04 | 0.43 | -0.60 | 0.88 |
| 16S-ITS-23S rRNA operon | -0.15 | -0.17 | 0.13 | -0.34 | 0.14 |

Table S7. Raw read count and consensus read count of each sample.

| **Sample ID** | **Raw Read Count** | **Consensus Read Count** |
| --- | --- | --- |
| **Glycerol_1** | 86,951 | 1,177 |
| **Glycerol_2** | 126,696 | 1,169 |
| **Glycerol_3** | 199,844 | 2,497 |
| **Glycerol_4** | 252,838 | 1,932 |
| **Glycerol_5** | 147,744 | 1,364 |
| **Glycerol_6** | 190,174 | 1,505 |
| **PCE_1** | 62,392 | 578 |
| **PCE_2** | 106,482 | 1,497 |
| **PCE_3** | 100,621 | 994 |
| **PCE_4** | 69,468 | 996 |
